# Natural Killer activating multimeric immunotherapeutic complexes (NaMiX) induce cytotoxic activity and killing of HIV-1 infected cells

**DOI:** 10.1101/2022.11.11.516216

**Authors:** Rafaela Schober, Bianca Brandus, Thessa Laeremans, Gilles Iserentant, Géraldine Dessilly, Jacques Zimmer, Michel Moutschen, Joeri L Aerts, Xavier Dervillez, Carole Seguin-Devaux

## Abstract

HIV-1 persists in viral reservoirs of latently infected CD4^+^ T cells containing integrated replication-competent viral DNA. Combined Antiretroviral Therapy (cART) does not eradicate HIV-1 reservoirs and treatment interruption will ultimately lead to viral load rebound. HIV-1 infection dramatically reduces the proportion of functional NK cell subsets and increases the expression of the checkpoint inhibitors NKG2A and KIR2DL. In this regard, we developed novel recombinant molecules combining multimers of the IL-15/IL-15Rα complex with the single-chain fragment variables (scFvs) of NKG2A or KIR2DL, and named them as Natural killer activating Multimeric immunotherapeutic compleXes (NaMiX). NaMiX significantly improved the cytotoxic activity of NK cells against HIV-1 positive ACH-2 cells and resistant Raji cancer cells by increasing their degranulation capacity, release of granzyme B, perforin and IFN-γ expression. Targeting the NKG2A receptor had a stronger effect compared to the targeting of the KIR2DL receptor due to its higher expression on NK cells. In a viral inhibition assay using CD4^+^ T cells from HIV-1 positive patients under cART, NaMiX initially increased viral replication which was subsequently inhibited by stimulated NK cells. In humanized NSG tg-huIL-15 mice showing functional NK cells, we observed enhanced activation, degranulation and killing by NK cells from the spleen of mice treated with anti-NKG2A NaMiX compared to the cells of control mice previously infected with HIV-1 and treated with cART. Although NaMiX did not delay viral load rebound after treatment interruption in a first attempt, it tend to decrease total HIV-1 DNA in the lungs of the mice. Blocking the inhibitory receptor NKG2A in combination with targeted multimers of IL-15 on NK cells could therefore be a promising immunotherapeutic strategy towards HIV-1 functional cure.

## Introduction

Combined Antiretroviral Therapy (cART) has transformed HIV-1 infection from a lethal disease into a chronic, manageable infection, considerably improving survival and prevention of transmission. However, HIV-1 persists in reservoirs of latently infected CD4^+^ T cells containing a minor part of integrated replication-competent viral DNA that is transcriptionally silent [1]. Given both the 38 million people living with HIV-1 worldwide and the inability of cART to eradicate the viral reservoir, the HIV epidemic remains one of the greatest health challenges in modern history. This underlines the urgency to find a cure. Two approaches have been investigated in this respect: on the one hand, HIV-1 research has during the last decade explored ways for the complete eradication of the reservoir, and on the other hand, considered a long-term remission approach in the absence of cART called “functional cure”. Although this terminology has been controverted [2], the concept relies on a cART-free ‘durable virologic suppression’ without clearing the latent reservoirs. The most studied reservoir eradication strategy is the “shock and kill” therapy, which relies on the activation of latent reservoir cells by Latency Reversal Agents (LRAs) followed by the recognition and elimination of cells harboring the reactivated virus by Natural Killer (NK) cells and cytotoxic T lymphocytes (CTL).

NK cell-based immunotherapy is a promising field in cancer treatment and is progressively proposed as an approach to control HIV-1 infection [3]. NK cells are multipotent innate lymphoid cells that have a fundamental function in immune-surveillance against cancer cells and virus infected cells without prior stimulation. They can rapidly recognize stressed cells through germline encoded activating or inhibitory NK cell receptors (aNKRs and iNKRs, respectively) and eliminate them through Antibody-Dependent Cellular Cytotoxicity (ADCC) mediated by the FcγRIIIA receptor (CD16) or exocytosis of lytic granules containing granzymes and perforin. Human NK cells represent 5-15% of circulating lymphocytes and make an important link to the adaptive immune system through the secretion of cytokines and chemokines and further interaction with dendritic cells. Two main NK cell populations are subdivided by their CD56 expression; the regulatory CD56^bright^ (5-15% of all peripheral blood NK cells), and the cytotoxic CD56^dim^ (90% of all peripheral blood NK cells). While NK cells are one of the first responders during the acute phase of HIV-1 infection and produce a large amount of IFN-γ [4], chronic infection generates abnormal distribution of subpopulations with the expansion of dysfunctional CD56^neg^ cells [5, 6], reduction of the aNKRs expression [7], down-regulation of cytokine production and reduction of stored perforin and granzyme A. cART is only partially able to recover NK cell distribution, cytotoxicity and IFN-γ expression [8, 9], which likely prevents them from clearing the latent reservoir after viral reactivation [10].

The ability of interleukin-15 (IL-15) to increase NK and T cell activation, expansion and proliferation has been well-established [11, 12] and novel IL-15 based therapies were extensively tested in oncology [13–16]. IL-15 is a regulatory cytokine from the common γ chain family and is predominantly trans-presented by antigen presenting cells (APCs) or cis-presented to target cells by its co-receptor unit α (IL-15Rα) [17]. The concentration of IL-15 must be constantly above a certain threshold to reach an effect on NK cell expansion and activation. However, IL-15 has a very short half-life (2.5h) making the *in vivo* administration of a single injection impossible without a high C_max_ and resultant toxicity [18]. Complexing IL-15 to its co-receptor IL-15Rα has proven to increase stability, solubility and even stimulatory activity [19]. IL-15/IL-15Rα based immunomodulatory molecules have shown a high potency to activate and increase cytotoxic activity of NK cells in the case of HIV-1 infection [20–23]. In addition, the superagonist ALT-803 (also called N-803), is able to activate latently infected cells, prime resting CD4^+^ T cells for CD8^+^ T cell recognition *in vitro* and *ex vivo,* and was proposed as a LRA [24]. Several clinical trials were started to assess the effect of ALT-803 on the control of HIV-1 infection in humans as a functional cure (NCT04505501), on acute infection (NCT04505501) or as a LRA (NCT04808908).

Blockade of inhibitory receptors such as cytotoxic T-lymphocyte associated protein 4 (CTLA-4) or the Programmed cell death 1 (PD-1)/PD-L1 axis on T lymphocytes is another major strategy applied in cancer immunotherapy [25]. These treatments can induce strong responses, but only in a minority of patients. The activation of NK cells depends on the stimulation balance between aNKRs and iNKRs, making them excellent candidates for immune checkpoint blockade. Potential targets on NK cells are the Killer cell immunoglobulin-like receptors 2DL (KIR2DL) or the lectin-like receptor NKG2A. They are inhibitory receptors for Human Leukocyte Antigen class I-C (HLA-C) and α-chain-E (HLA-E) respectively. HLA-C and HLA-E are broadly expressed on healthy tissues to define immune “self”, but HLA-E can be over-expressed on cancer cells or HIV-1 infected T lymphocytes in order to escape immune recognition [26, 27]. Hence, down-regulation of NKG2A on NK cells increases antitumor activity against HLA-E expressing resistant tumor cells [28] and blocking NKG2A in combination with the PD-1/PD-L1 pathway improves tumor control [29]. Moreover, blocking the KIR2DL1, DL2, DL3 receptors increased NK cell mediated killing of acute myeloid leukemia cells and activated NK cells from HIV-1 infected viremic and aviremic patients [30].

In this study, we describe the development and the validation of novel therapeutic molecules, called NaMiX, for NK activating Multimeric immunotherapeutic compleXes. The oligomerization domain of C4 binding protein (C4bp) was used to associate the IL-15/IL-15Rα complex with an anti-NKG2A or an anti-KIR single-chain fragment variable (scFv) as well as to multimerize each entity into a heptamer (α form) or a dimer (β form) [31]. The different molecules were tested for their ability to activate NK cells and increase their killing against HIV-1-infected and cancer cells *in vitro*. Preliminary *ex vivo* and *in vivo* experiments were performed in humanized NSG tg-huIL-15 mice harboring functional NK cells to evaluate the lead anti NKG2A.IL-15 NaMiX.

## Results

### Molecular characterization

C4bp is a protein complex of seven identical α chains and a single β chain, which inhibits the lectin and classical pathways of the complement system. Each subunit is composed of several complement control proteins (CCPs) and an oligomerizing C-terminal domain. The inactive oligomerizing entities of C4bpα and C4bpβ have the unique ability of forming heptamers and dimers, respectively, by making disulfide bounds (Figure 1A). We previously reported that the fusion of effector or target entities into this domain does not affect its oligomerizing capacity and can be used to increase half-life [31] and functionality of chosen proteins [32]. To express IL-15 heptamers (α molecules) or dimers (β molecules) at the surface of NK cells and generate NaMiX, we grafted the extracellular sushi domain of IL-15Rα on the C-terminus of the C4bp oligomerization domain and the scFvs of anti-NKG2A or anti-KIR on the N-terminus. All molecule-coding sequences were cloned into pEF-IRES-pac expression vectors containing a histidine (His) tag and were co-transfected into HEK293F cells transfected with a rhuIL-15 coding pcDNA3.1 to produce α.anti-NKG2A.IL-15, β.anti-NKG2A.IL-15, α.anti-KIR.IL-15 and β.anti-KIR.IL-15 NaMiX. Control molecules without IL-15 were also generated for each anti-NKG2A or anti-KIR NaMiX. However, the only control molecule that was sufficiently produced without co-transfection with rhuIL-15, was β.anti-NKG2A, so the controls α.anti-NKG2A, α.anti-KIR and β.anti-KIR could not be considered. We first assessed the binding efficiency of IL-15 to its co-receptor IL-15Rα, using an anti-IL-15/anti-His sandwich ELISA on increasing concentrations of the NaMiXs (Figure 1B). The β.anti-NKG2A.IL-15 and β.anti-KIR.IL-15 forms required 0.26 and 0.32 μg of each NaMiX, respectively, to reach a 50% saturation, while the α.anti-NKG2A.IL-15 and α.anti-KIR.IL-15 forms only required 0.07 and 0.06 μg of each NaMix, respectively. Full saturation was reached at 0.375 μg for the α form and at 1.5 μg for the β forms. The α forms of the molecules have indeed seven His tags and IL-15 entities whereas the β forms only have two entities of each, explaining the 4-fold higher concentration needed to reach a plateau. Next, we wanted to determine if all IL-15Rα sites were saturated with IL-15 and whether external rhuIL-15 could still bound to NaMiX. When adding external rhuIL-15 to saturating concentrations of β.anti-NKG2A, it required 7.8 ng of rhuIL-15 to saturate 6 μg of the molecule indicating that 3 μg of the β.anti-NKG2A.IL-15 NaMiX used in this study is approximatively equivalent to 3.9 ng of rhuIL-15. As shown in Figure 1C, all sites of α.anti-NKG2A.IL-15 were saturated with IL-15 while α.anti-KIR.IL-15 required an additional 122 pg rhuIL-15 and β.anti-NKG2A.IL-15, and β.anti-KIR.IL-15 required an additional 0.5 μg rhuIL-15 to saturate all sites of the molecules.

**Figure 1:**
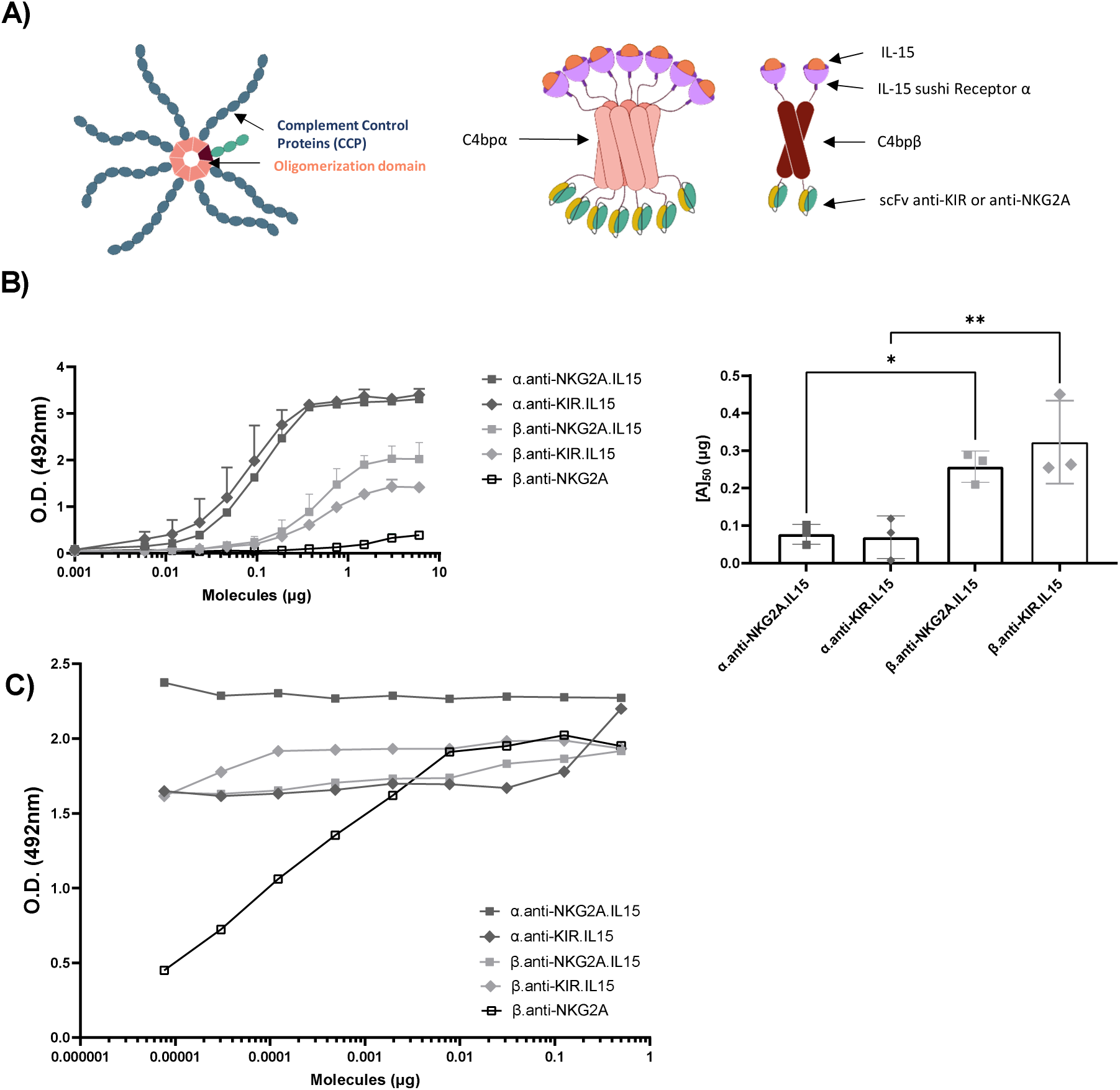
Demonstration of the huIL-15/ IL-15Rα complex formation on purified molecules. **A)** Schematic representation of the multimeric molecules. The alpha forms multimerize in heptamers while the beta forms multimerize in dimers. The anti-NKG2A or anti-KIR scFvs are located in N-terminal of the C4bp moiety while IL15Rα is located in C-terminal. The plasmids coding for the molecules are co-transfected with a plasmid coding for recombinant human IL-15. **B)** Polystyrene MaxiSorp™ plates were coated with a mouse anti-human IL-15 mAb, incubated with a two-fold serial dilution of molecules (6 μg to 0.001 μg) and binding of the molecules was detected with a mouse anti-HIS pAb HRP-conjugated. Figures 1B represent triplicates of three independent experiments; data are expressed as the mean value ± SD. The right panel represents the concentration of the NaMiX needed to half saturate the ELISA (left). Statistical analysis was performed using a one-way ANOVA and post-hoc Tukey test (* p < 0.05, ** p < 0.005). **C)** Saturating concentration of molecules were coated on Polystyrene MaxiSorp™ plates (6 μg for β forms and 3 μg for α forms). A serial dilution of rhuIL-15 was added to each molecule. The binding of IL-15 was detected with a mouse anti-human IL-15 mAb followed by a goat anti-Mouse IgG HRP-conjugated pAb. Figure 1C shows the data of one representative experiment.

### NaMiX bind to their respective receptors NKG2A and KIR2DL1/2DL2/2DL3

We first confirmed that all NaMiX bound to their respective receptors using KIR2DL1, 2DL2, 2DL3-expressing HEK293F cells and on NKG2A-expressing NK-92MI cells by flow cytometry (Figure2A). Both α.anti-KIR.IL-15 and β.anti-KIR.IL-15 were found to bind to all target KIR2DL receptors expressed on stable cell lines. Similarly, α.anti-NKG2A.IL-15, β.anti-NKG2A.IL-15 and the β.anti-NKG2A control molecule, bound all to the NKG2A expressing NK-92MI cell line (Figure 2B). Since the NKG2A and KIR receptors are heterogeneously expressed on human NK and CD8^+^ T cells, we measured their expression on Peripheral Blood Mononuclear Cells (PBMCs) from healthy donors (Figure 2C) and evaluated the subsequent binding of NaMiX on these cells. NKG2A was expressed on more than 50% of the CD3-CD56+CD16+ NK cells whereas the total expression of all KIRs was not exceeding 35% of PBMC-derived NK cells. As shown by the His staining, the α.anti-NKG2A.IL-15 NaMiX bound to 83.7% ± 4.041% of all NK cells while the β.anti-NKG2A.IL-15 form only bound to 20.7% ± 11.21% of NK cells (figure 2D), in agreement with the respective number of anti-NKG2A scFv valences of the α heptamer and the β dimer. When looking at the IL-15 positive NK cells, although the reduced binding between the β and α form of the NKG2A NaMiX was significant (p=0.0006), α.anti-KIR.IL-15 did not show a higher signal than β.anti-KIR.IL-15 (p=0.998) suggesting a lower IL-15 amount on α.anti-KIR.IL-15 (Figure 1C) than α.anti-NKG2A.IL-15, in accordance with data from Figure 1C.

**Figure 2:**
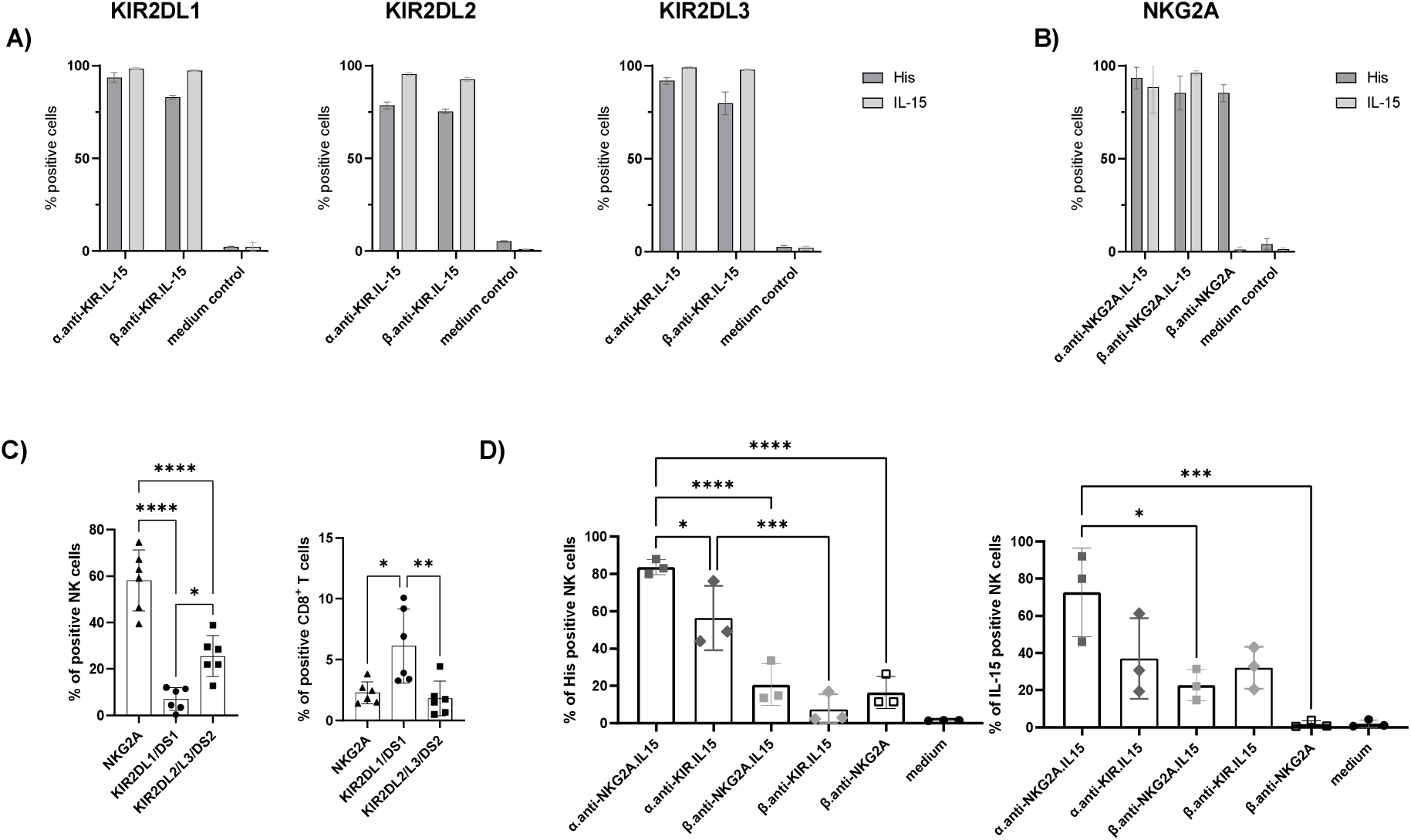
NaMiX bind to their respective receptors on NK cells. **A)** The molecules containing the anti-KIR scFv were incubated during 30 minutes with HEK293F cells expressing KIR2DL1 (left), KIR2DL2 (middle) and KIR2DL3 (right) and stained with anti-His and anti-IL-15 for flow cytometry analysis. **B)** The molecules containing the anti-NKG2A scFv were incubated with the stable cell line NK-92MI expressing NKG2A and stained with anti-His and anti-IL-15 for flow cytometry analysis. **C)** PBMCs from different donors were stained for extracellular markers including KIR2DL1/DS1, KIR2DL2/L3/DS2 and NKG2A to identify and phenotype CD3^-^CD56^+^CD16^+^ NK cells (left) and live CD3^+^CD8^+^ T (right) cells using anti-CD3, CD8, CD14, CD16, CD19 and CD56 antibodies **D)** All NaMiXs were incubated with human PBMCs and stained for NK cell markers, anti-His and anti-IL-15 for flow cytometry analysis. Data were expressed as the mean value ± SD. Figure A and B represent three independent experiments with tree different donors. Figure C and D represents six and three independent experiments, respectively, with three and six different donors. Statistical analysis was performed using a one-way ANOVA and post-hoc Tukey test (*p < 0.05, **p < 0.005, ***p < 0.0005, ****p < 0.00005).

### NaMiX increased higher STAT5 phosphorylation in NK and CD8^+^ T cells than recombinant human IL-15

Upon binding to IL-15, the cytokine receptors complex recruits the tyrosine kinases JAK1 and JAK3 to phosphorylate STAT5 (pSTAT5) and induce signaling pathways leading to cell survival [33, 34]. We investigated the JAK/STAT5 phosphorylation pathway on PBMCs by intracellular staining using flow cytometry and multispectral imaging cytometry. We first confirmed the intracellular STAT5 phosphorylation by imaging flow cytometry (Figure 3A) and observed that all molecules containing IL-15 definitely induce pSTAT5 signal on NK cells and CD8^+^ T cells, whereas β.anti-NKG2A without IL-15 and medium control showed no signal. We further observed an overall strong increase in pSTAT5 positive NK and CD8^+^ T cells early after incubation with α.anti-NKG2A.IL-15 compared to rhuIL-15 alone and β.anti-NKG2A.IL-15 (Figure 3A). Finally, STAT5 was less phosphorylated by anti-KIR.IL-15 NaMiXs than α.anti-NKG2A.IL-15 in NK cells and the β.anti-NKG2A control molecule did not induce any STAT5 phosphorylation.

**Figure 3:**
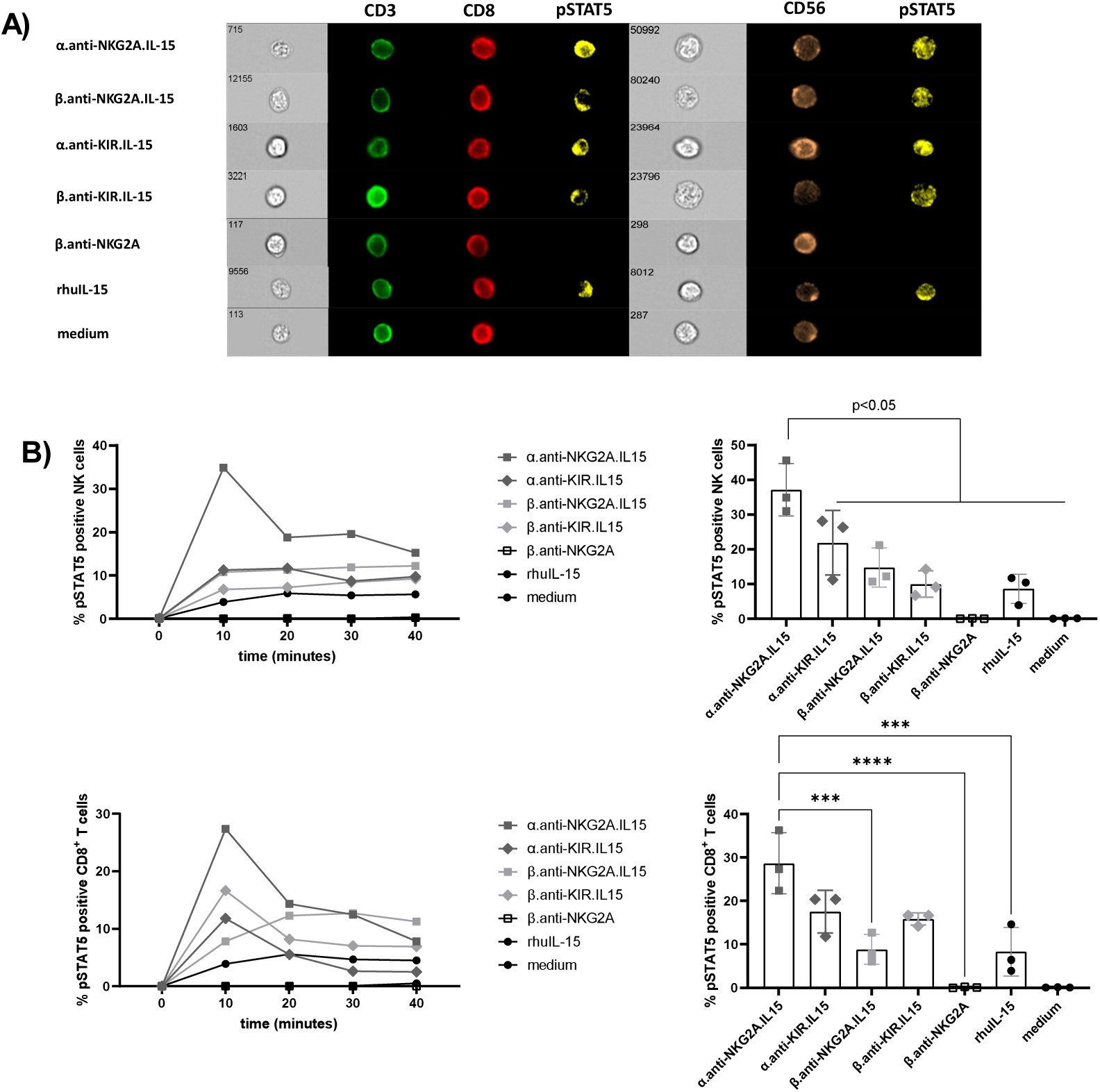
α.anti-NKG2A.IL-15 increased STAT5 phosphorylation in NK and CD8+ T cells. **A)** PBMCs were incubated with the indicated molecules for 1 minute, stained on ice to gate for CD3^-^CD56^+^CD16^+^ NK cells and CD3^+^CD8^+^ T cells using anti-CD3, CD8, CD14, CD16, CD19 and CD56 antibodies, permeabilized on ice and stained for intra-cellular pSTAT5 for imaging flow cytometry. **B)** PBMCs were incubated with the NaMiXs molecules for 1, 10, 20 or 40 minutes, stained on ice to gate for live CD3^-^CD56^+^CD16^+^ NK (upper panel) and live CD3^+^CD8^+^ T cells (lower panel), permeabilized on ice and stained for intra-cellular pSTAT5 for flow cytometry analysis. The left panels represents time-dependent pSTAT5 phosphorylation of one representative donor. Right panel represents three different healthy donors at 1 min incubation. Data were expressed as the mean value ± SD. Statistical analysis was performed using a one-way ANOVA and post-hoc Tukey test (***p < 0.0005, ****p < 0.00005).

### NaMiX significantly increased NK cell degranulation and cytotoxic activity against Raji cells

We first evaluated the effect of NaMiXs on PBMCs regarding IFN-γ and TNF-α secretion after 48h stimulation. All molecules except β.anti-NKG2A resulted in a significantly increased IFN-γ and TNF-α secretion as compared to medium control or treatment with rhuIL-15. However, α.anti-NKG2A.IL-15 and β.anti-NKG2A.IL-15 had a stronger effect than α.anti-KIR.IL-15 and β.anti-KIR.IL-15 on IFN-γ secretion, respectively (p<0.01) (Figure 4). The α.anti-NKG2A.IL-15 NaMix had a significantly stronger effect than α.anti-KIR.IL-15 on TNF-α secretion (p=0.0177) but this was not the case for the β forms. Furthermore, β.anti-NKG2A without IL-15 did not stimulate IFN-γ or TNF-α secretion suggesting that blocking of the target receptors with the anti-NKG2A ScFv did not induce cytokine secretions. Regarding granzyme B and perforin, their degranulation was significantly increased compared to medium control and rhuIL-15. Taken together, these data indicate that PBMCs were stimulated by NaMiXs to degranulate and produce cytokines even in the absence of target cells.

**Figure 4:**
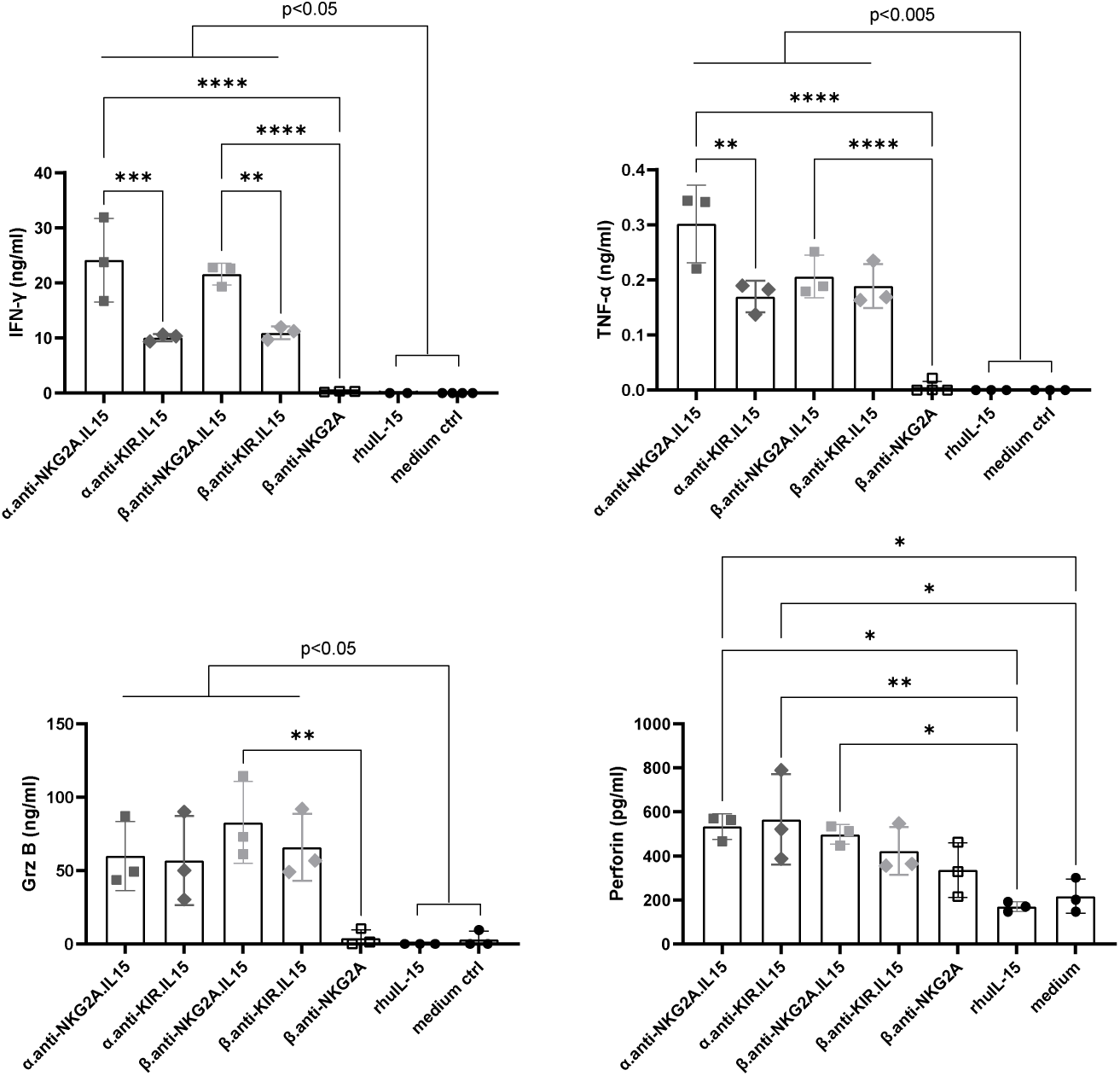
NaMiX increased IFN-γ and TNF-α secretion of PBMCs in vitro. Human PBMCs were incubated for 48h with the different NaMiXs. The supernatant was tested for the presence of IFN-γ (upper left panel), TNF-α (upper right panel), granzyme B (lower left panel) and perforin (lower right panel) by ELISA. The figure represents three independent experiments with three different healthy donors. Data were expressed as the mean value ± SD. Statistical analysis was performed using a one-way ANOVA and post-hoc Tukey test (**p < 0.005, ***p < 0.0005, ****p < 0.00005).

The next step was to evaluate the capacity of NaMiXs to induce NK cell degranulation and cytotoxicity against Raji cancer cells using PBMCs stimulated with the different molecules for 24 and 48h. Raji is a B-lymphoma cell line constitutively expressing CD20 and overexpressing multiple HLA class-I molecules, making it resistant to NK cell recognition and killing in the absence of additional stimuli. Pre-stimulation of PBMCs for 48h with α.anti-NKG2A.IL-15 significantly increased surface expression of the degranulation marker CD107a on NK (mean 76.25% ± 10.89%) and CD8^+^ T cells (mean 75.22% ± 6.07%) compared to rhuIL-15 (p=0.0405; p=0.0010, respectively) and medium control (p= 0.0036; p= 0.0001, respectively) (Figure 5A) against Raji cells whereas the anti-KIR.IL15 molecules had no significant effect. Furthermore, anti-KIR.IL-15 did not affect CD107a expression of CD8^+^ T cells despite a slightly higher KIR2DL1/DS1 receptor expression compared to NKG2A receptors as shown in Figure 2C. Importantly, 24h stimulation with α.anti-NKG2A.IL-15 and β.anti-NKG2A.IL-15 molecules improved the ability of NK cells to express IFN-γ (mean 29.09% ± 10.61%; mean 31.68% ± 11.23%, respectively) against Raji target cells compared to rhuIL-15 stimulation alone (p= 0.028; p=0.014 respectively) (Figure 5B), which was not observed in CD8^+^ T cells (Figure 5C). Moreover, β.anti-NKG2A stimulation showed no increase in IFN-γ secretion compared to medium control (Figure 5A and B) indicating that the effect of the molecules on IFN-γ expression and secretion originates from IL-15 stimulation rather than the blocking of NKG2A. The α.anti-NKG2A.IL-15 was superior only in CD8^+^ T cells to increase CD107a after 48 hours of stimulation as compared to the other molecules, but this was not the case in NK cells, and was not in accordance with the expression/secretion of IFN-γ, granzyme B and perforin. When looking at the granzyme B secretion, we only observed a significant increase when PBMCs were stimulated with the NKG2A NaMiX (both α- and β-based NaMiXs) compared to rhuIL-15 and medium control, whereas perforin was increased with all molecules except with the β.anti-NKG2A control. In summary, these results suggest that both α.anti-NKG2A.IL-15 and β.anti-NKG2A.IL-15 have a strong potency on increasing cytotoxic activity of NK and CD8^+^ T cells against resistant cancer cells.

**Figure 5:**
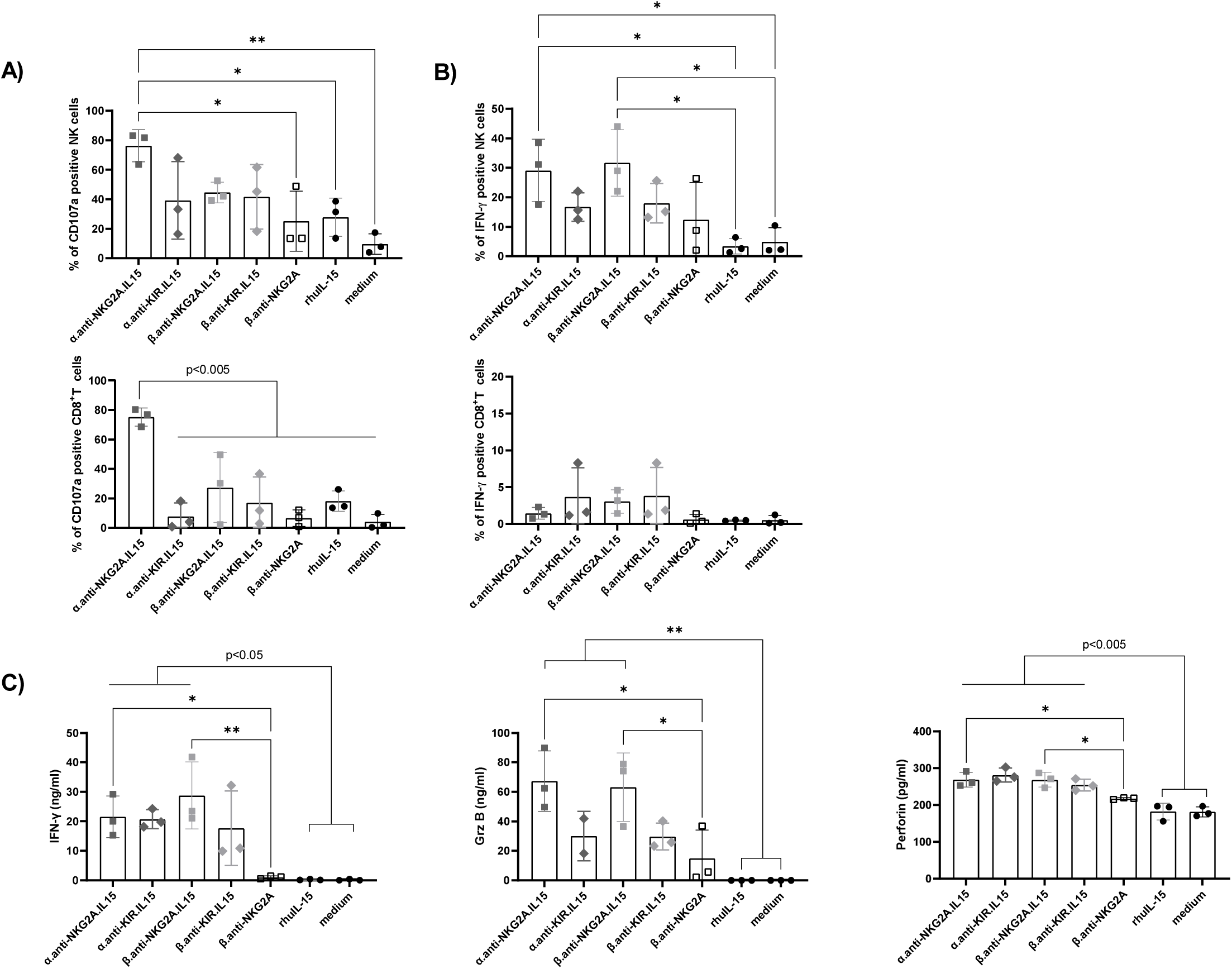
NaMiX increased degranulation and IFN-γ secretion of NK cells against resistant Raji cells in vitro. **A)** Human PBMCs were pre-incubated for 24h (for IFN-γ expression) or 48h (for CD107a expression) with the NaMiX and stimulated with Raji cells for 5h in presence of anti-CD107a mAb. Cells were further stained for extracellular markers to identify CD107a expression on CD3^-^CD56^+^CD16^+^ NK cells (upper panel) and CD3^+^CD8^+^ T cells (lower panel) using anti-CD3, CD8, CD14, CD16, CD19 and CD56 antibodies. **B)** Cells were also permeabilized and further stained for IFN-γ (NK cells on upper panel and CD8^+^ T cells on lower panel). **C)** After incubation, supernatant was collected and analysed by ELISA for IFN-γ (left panel), granzyme B (middle panel) and perforin secretion (right panel). Figures 5A to 5C represent three independent experiments. Data were expressed as the mean value ± SD. Statistical analysis was performed using a one-way ANOVA and post-hoc Tukey test (*p < 0.05, **p < 0.005).

### NaMiX significantly increased NK cell degranulation and cytotoxic activity against activated ACH-2 cells

We further tested the effect of the molecules on NK and CD8^+^ T cell degranulation and cytotoxicity against activated ACH-2 cells, an acute lymphoblastic leukemia T-cell line containing a single integrated copy of HIV-1 per cell. Viral expression in ACH-2 cells was induced by PHA/PMA mediated cell activation [35], which resulted in high intracellular p24 expression (data not shown). 48h stimulation with NaMiXs had a greater effect on CD107a surface expression of NK cells and CD8^+^ T cells against HIV-1 positive cells than rhuIL-15 alone (Figure 6A). Interestingly, β.anti-NKG2A without IL-15 (mean 60.7% ± 6.47%) had a similar effect than the other molecules, suggesting that blocking the NKG2A receptor in the context of HIV-1 infection had a more robust effect on degranulation as compared to Raji cells. The secretion of IFN-γ by NK cells was similarly increased by all IL-15 NaMiX (except for β.anti-NKG2A) compared to rhuIL-15 and medium control. Surprisingly, all molecules, even β.anti-NKG2A without IL-15 also increased IFN-γ secretion by CD8^+^ T cells (Figure 6B). This indicates that blocking the NKG2A pathway is sufficient to enhance IFN-γ production by CD8^+^ T cells but not by NK cells. These data were further confirmed by measuring IFN-γ, granzyme B and perforin secretion in the supernatant (Figure 1 C). All NaMiXs showed enhanced IFN-γ, granzyme B and perforin secretion into the supernatant but not the β.anti-NKG2A control compared to rhuIL-15 and medium control (Figure 6C). To evaluate the potential involvement of ADCC, we also performed the experiment in the presence of sera from HIV-1 positive and negative donors, respectively, and observed no differences in the degranulation capacity or IFN-γ expression of NK and CD8^+^ T cells between both conditions (data not shown). This indicates that the increase in degranulation and cytokine release by NaMix is independent of ADCC and relies only on direct cytolytic activity. Taken together, all these data confirmed that all NaMiX molecules were able to stimulate NK and CD8^+^ T cells *in vitro* against an HIV-infected cell line.

**Figure 6:**
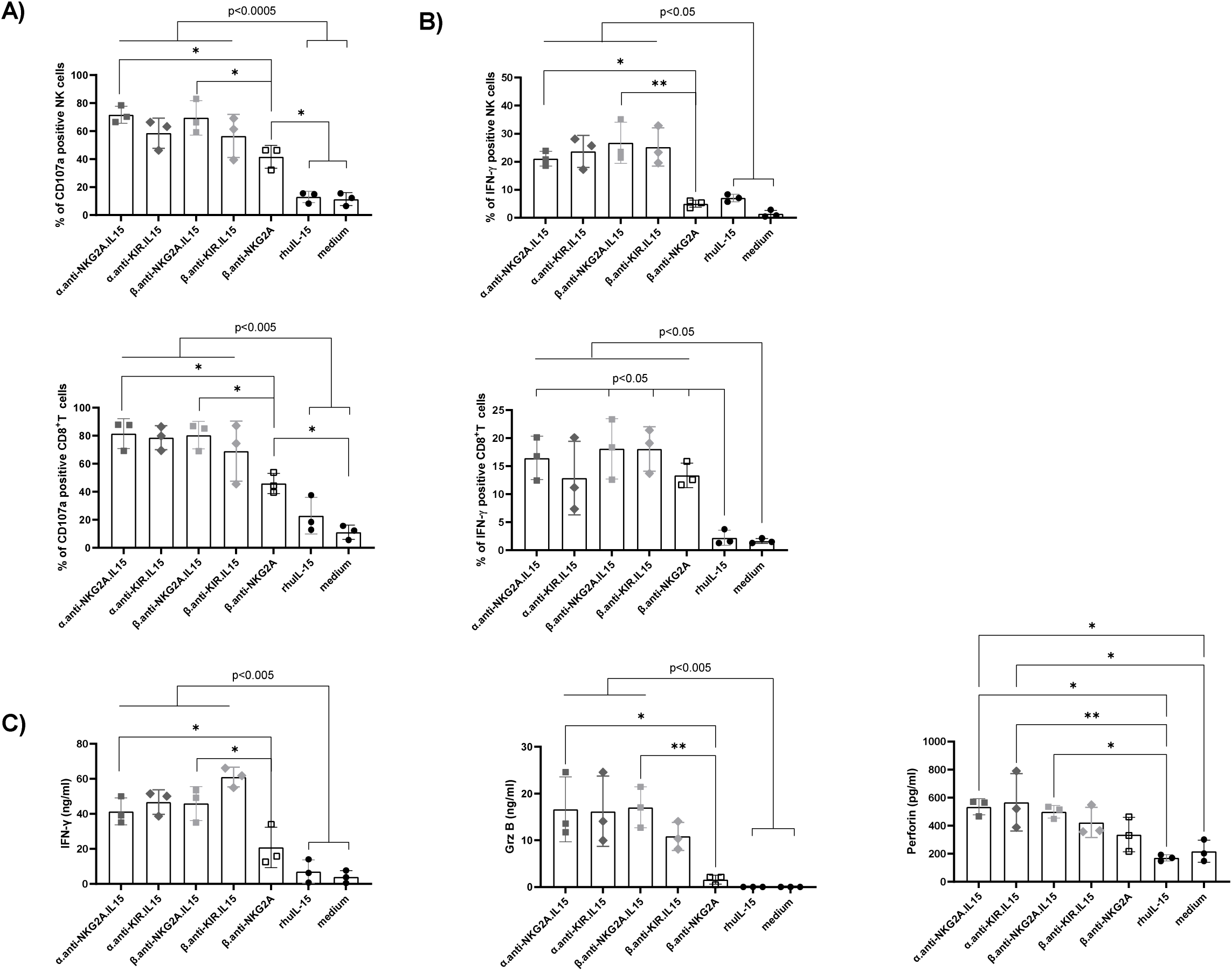
NaMiX increased degranulation and IFN-γ secretion of NK and CD8^+^ T cells against activated ACH-2 cells in vitro. **A)** Human PBMCs were pre-incubated for 48h with the different molecules and stimulated with the activated ACH-2 cells for 5h in presence of anti-CD107a. Cells were further stained for extracellular markers to identify CD107a expression on CD3^-^CD56^+^CD16^+^ NK cells (upper panel) and CD3^+^CD8^+^ T cells (lower panel) using anti-CD3, CD8, CD14, CD16, CD19 and CD56 antibodies. **B)** Cells were also permeabilized and further stained for IFN-γ (NK cells on upper panel and CD8^+^ T cells on lower panel). **C)** After incubation, supernatant was collected and analyzed by ELISA for IFN-γ (left panel), granzyme B (middle panel) and perforin secretion (right panel). Figures 6A to 6C represent three independent experiments. Data were expressed as the mean value ± SD. Statistical analysis was performed using a one-way ANOVA and post-hoc Tukey test (*p < 0.05, **p < 0.005).

### NaMiX significantly enhanced NK cell cytotoxic killing of Raji and activated ACH-2 cells

We next evaluated the killing capacity of NK and CD8^+^ T cells pre-stimulated for 48 hours with NaMiX against the targets Raji or ACH-2. In Figure 7A, PBMCs pre-incubated with all IL-15 NaMiXs showed a strong killing activity ranging from 70% to 90% against Raji cells when measuring Live/Dead cells by flow cytometry while rhuIL-15 alone, β.anti-NKG2A and the medium control showed a killing activity of 38% (± 10.51%), 28% (± 6.9%) and 18% (± 3.16%), respectively. Furthermore, β.anti-NKG2A.IL-15 stimulated PBMCs killed 92% (± 3.62%) of Raji cells as compared to 28% (± 6.91%) for β.anti-NKG2A, emphasizing the requirement of IL-15 for an optimal cytotoxicity against resistant cancer cells. However, when measuring dehydrogenase activity as a marker of cell viability, (Figure 7B), α.anti-NKG2A.IL-15 and β.anti-NKG2A.IL-15 stimulation showed a significantly higher killing activity compared to either α or β.anti-KIR.IL-15, respectively (p= 0.036; and p=0.044, respectively).

**Figure 7:**
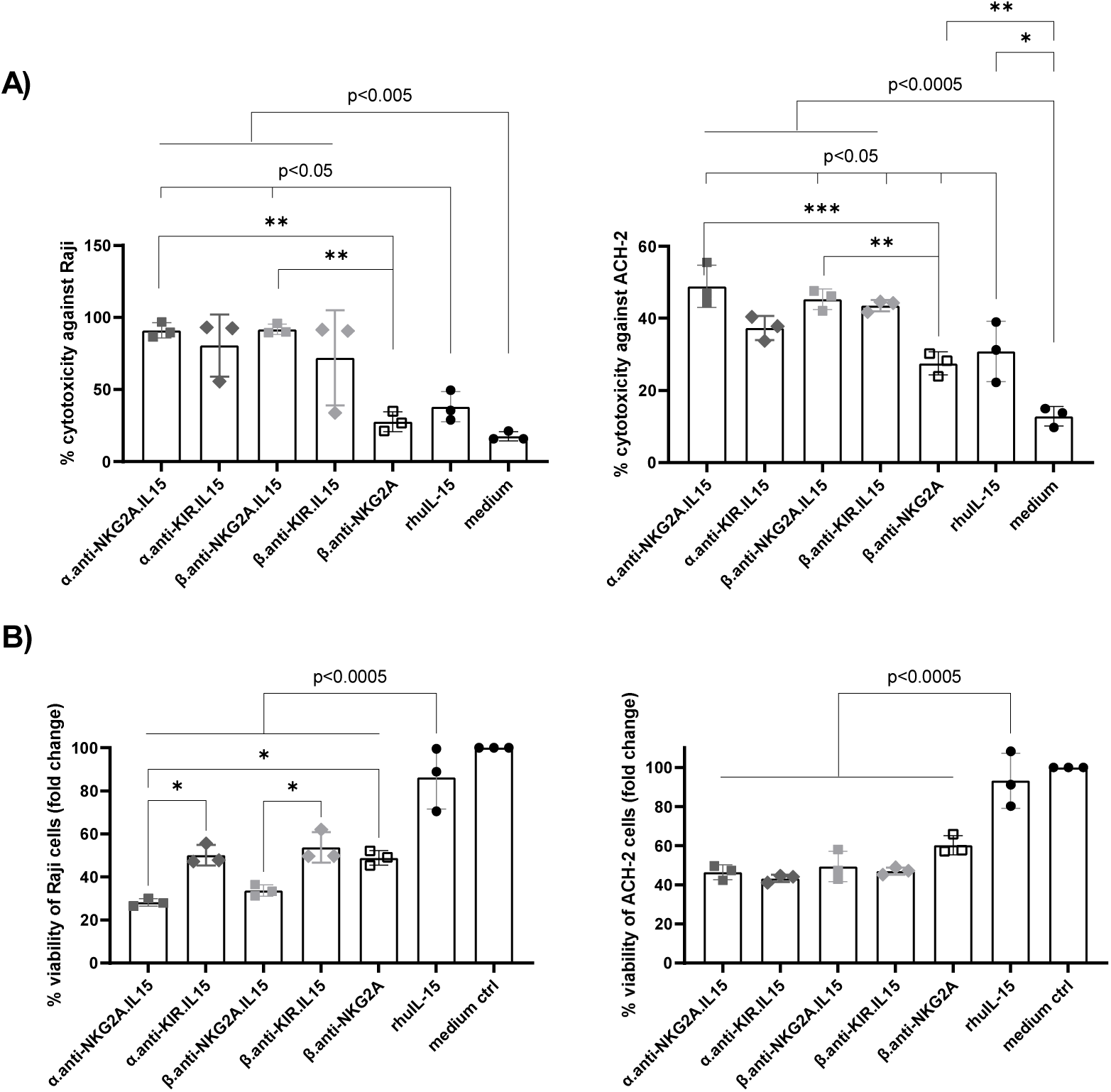
NaMiX enhanced the cytotoxic killing activity of NK and CD8^+^ T cells against Raji and activated ACH-2 cells in vitro. Human PBMCs were pre-incubated for 48h with the different molecules. **A)** Stimulated PBMCs were incubated with Raji (left panel) or ACH-2 (right panel) cells pre-stained with CellTrace Violet for 5h. Cells were further stained using Live/Dead staining and analyzed by flow cytometry. **B)** Stimulated PBMCs were incubated with Raji (left panel) or ACH-2 (right panel) cells for 5h and the CCK8 reagent was added after 2h. Absorbance was read at 450 nm. The figures represent independent experiments with three different donors. Data were expressed as the mean value ± SD. Statistical analysis was performed using a one-way ANOVA and post-hoc Tukey test (*p < 0.05, **p < 0.005, ***p < 0.0005).

When using activated ACH-2 cells, all NaMiX significantly increased the capacity of PBMCs to kill target cells to a similar level compared to rhuIL-15 alone (Figure 7B). When measuring the dehydrogenase activity in the supernatant, no differences were observed between the NaMiXs with a mean of 40% of viable cells. We also evaluated whether ADCC could be implicated in the killing of the target ACH-2 cells by adding sera from HIV-1 positive or negative donors, respectively, and did not observe any variation in the killing capacity (data not shown). Altogether, these data confirmed that the IL-15 NaMiXs increase the capacity of NK and CD8^+^ T cells to kill resistant cancer cells and HIV-1 positive target cell lines.

### IL-15 stimulation and blocking of NKG2A-HLA-E interaction are required for NK cell activation

Since we observed an increased activation and cytotoxicity of NK and CD8^+^ T cells stimulated with β.anti-NKG2A without IL-15 when co-incubated with ACH-2 cells, we wanted to evaluate further whether HLA-E blocking is involved in the effect of the anti-NKG2A NaMiXs. We therefore incubated PBMCs together with the anti-NKG2A NaMiXs and K562 cells modified to express only HLA-E and are devoid of all MHC molecules, and were. We observed that blocking HLA-E with β.anti-NKG2A alone was insufficient to establish neither CD107a surface appearance nor IFN-γ expression by NK cells compared to medium control (Figure 8A). However, β.anti-NKG2A.IL-15 significantly increased CD107a expression as compared to β.anti-NKG2A alone (p=0.0227). We did not observe the expected increase of degranulation by the α form as seen with the Raji cells, which could be due to the fact that the cells were not pre-stimulated during 48 h with NaMiX in this setting. Furthermore, only the α.anti-NKG2A.IL-15 was able to significantly increase IFN-γ expression of NK cells compared to medium control and rhuIL-15 (p=0.0481 and p=0.0117, respectively) (Figure 8B). Interestingly, even though α.anti-NKG2A.IL-15 increased the killing of HLA-E expressing K562 cells, only β.anti-NKG2A.IL-15 was able to show a significant increase compared to medium control and β.anti-NKG2A (p=0.0183 and p=0.0194, respectively) (Figure 8C). We did not observe any difference in the degranulation and IFN-γ expression of CD8^+^ T cells (data not shown). Overall, these results suggest that the administration of rhuIL-15 or blocking the interaction with HLA-E alone is not sufficient to increase the cytotoxic capacity of NK cells but rather requires the synergy of both mechanisms.

**Figure 8.**
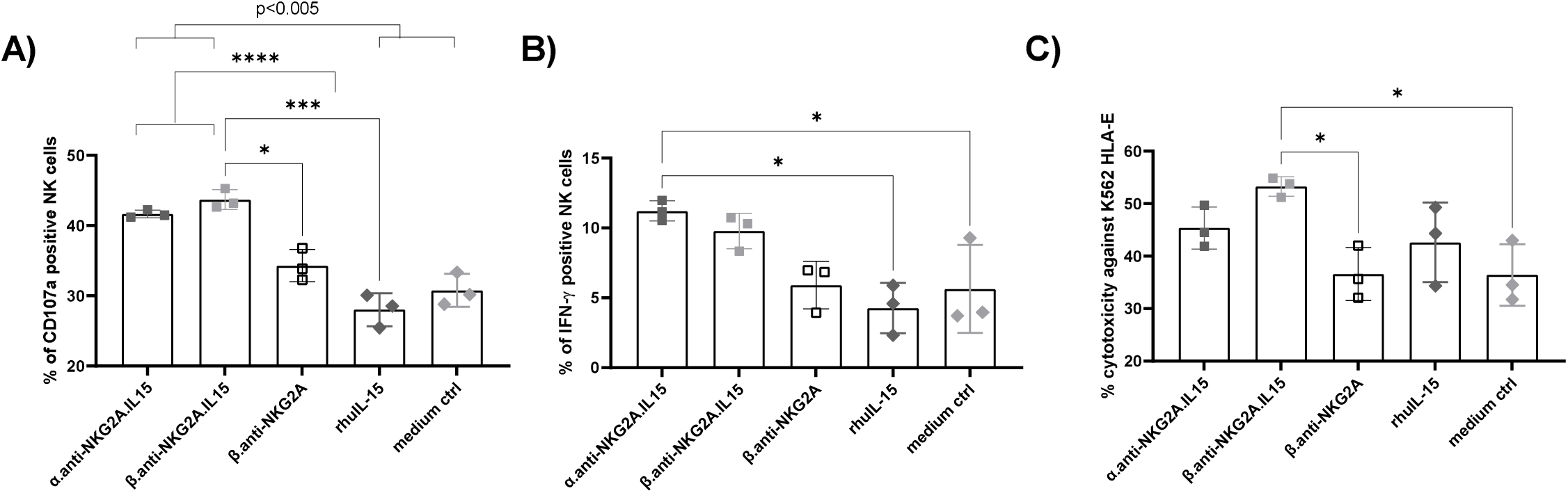
IL-15 stimulation and blocking of NKG2A-HLA-E interaction were required for NK cell activation by NKG2A NaMiX. **A)** Human PBMCs were incubated for 5h with the NKG2A NaMiX and HLA-E expressing K562 cells in presence of anti-CD107a mAb. Natural Killer cells were further stained for extracellular markers to identify CD107a expression. **B)** Cells were also permeabilized and further stained for IFN-γ. **C)** Human PBMCs were incubated for 5h with the NaMiX and HLA-E expressing K562 pre-stained with CellTrace Violet. Cells were further stained for Live/Dead and analyzed by flow cytometry. The figure represents three independent experiments with different donors. Data were expressed as the mean value ± SD. Statistical analysis was performed using a one-way ANOVA and post-hoc Tukey test (**p < 0.005, ***p < 0.0005, ****p < 0.00005).

### NaMiX reduced HIV-1 replication in a viral inhibition assay

We next examined the effect of NaMiXs on the viral suppressive capacity of NK cells of patients chronically infected with HIV-1 under cART. We observed increased p24 positive CD4^+^ T cells after two days of incubation with NK cell stimulated with all molecules and rhuIL-15 (Figure 9A) suggesting a higher replication of the virus when stimulated with NaMiXs at earlier time point that could be due to the activation of NK and T cells and the potential of IL-15 to reverse latency. Due to heterogeneous basal infection level, the mean of change of the different donors was very large, but the same tendency was observed in all three donors evaluated. After five days co-incubation with both α and β-IL-15-NaMiXs or rhuIL-15 stimulated NK cells, a decrease of p24 positive CD4^+^ T cells was further measured (Figure 9A and 9B). The β.anti-NKG2A control without IL-15 increased viral replication at two days co-incubation but did not decrease the intracellular p24 level after five days indicating that IL-15 alone is responsible of NK cell mediated viral inhibition. When looking at viral RNA in the supernatant, the levels were also slightly increased at two days co-incubation with stimulated NK cells compared to medium control (Figure 9B). In contrast, after five days of co-incubation with stimulated NK cells, NaMiX induced a decrease of HIV-1 RNA as compared to untreated NK cells, this effect was similar to rhuIL-15 for the α NaMiXs with IL-15. We further checked whether NaMiX was indeed inducing or decreasing viral replication by activating PBMCs. As shown in Figure 9B, the activation marker CD25 was significantly increased by all forms of NaMiX in NK cells (P < 0.00005) and in CD4+ T cells (P < 0.05) but not with β.anti-NKG2A whereas the CD69 marker was induced by all molecules in NK cells (P < 0.00005) but not in CD4+ T cells. Next, we assessed the release of HIV-1 mRNA after 24 hours of co-culture (Figure 9D) and observed an overall decrease of viral replication by all NaMiX as compared to non-stimulated ACH2 cells or IL-15, but that was not significant. Earlier time points showed low and similar mRNA levels among the different conditions, and the level of P24 expression measured by flow cytometry and ELISA was also too low to observe a significant effect with ACH2 cells at early time points (data not shown). Taken together, these results suggests that NaMiX decrease viral replication of HIV-1 infected cells after cellular activation and killing of CD4+ T cells by stimulated NK cells.

**Figure 9:**
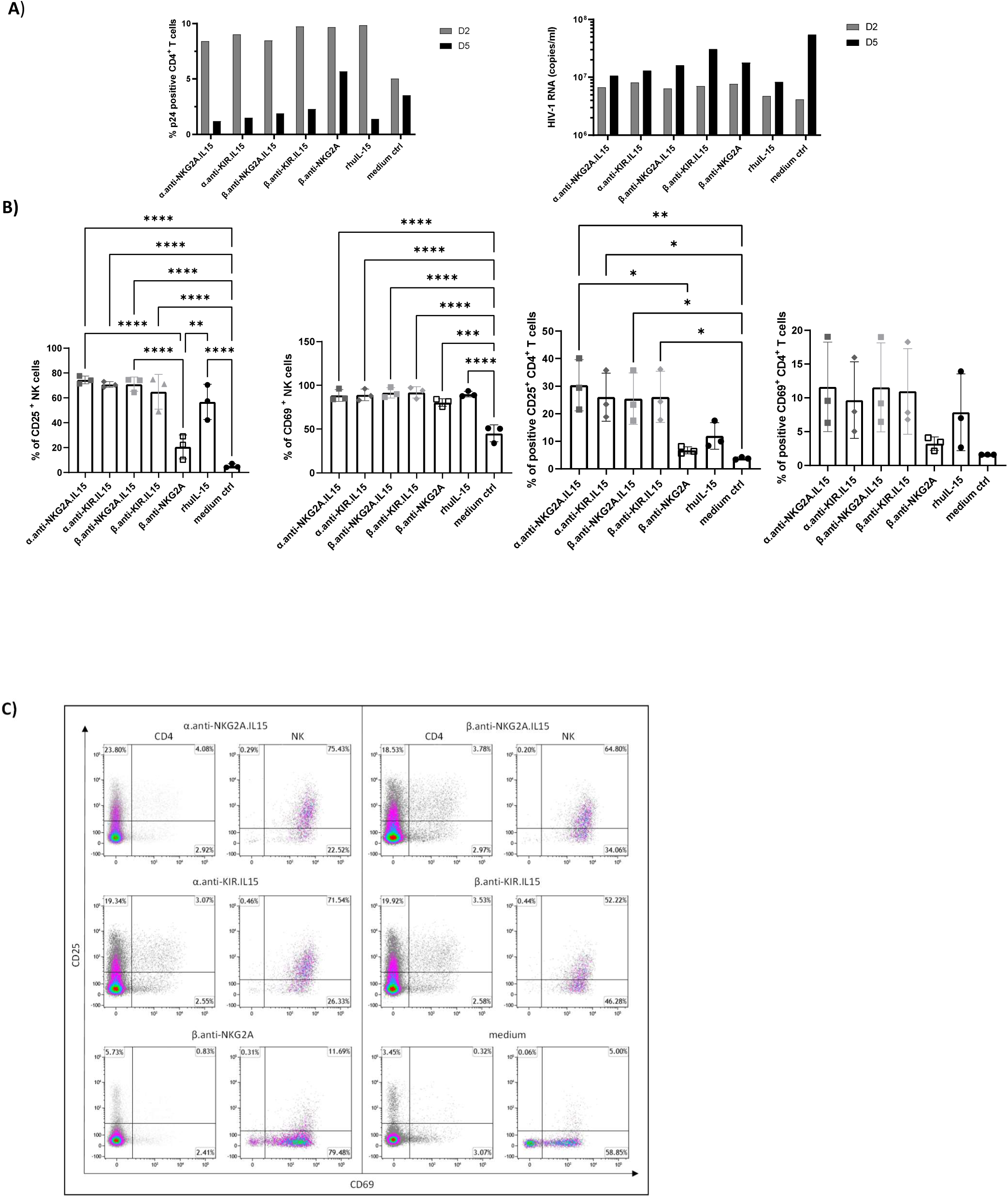
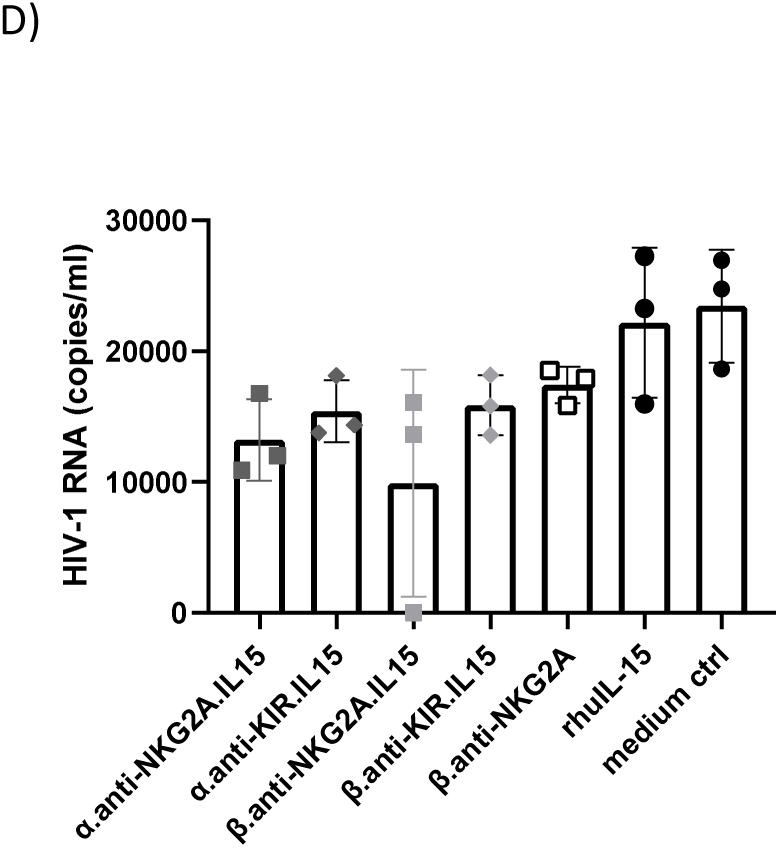
NaMiX increased viral inhibition capacity of NK cells against HIV superinfected CD4^+^ T cells from cART-treated patients by activating and killing CD4^+^ T cells. Autologous CD4^+^ T cells and NK cells were purified by positive and negative selection, respectively, from PBMCs of cART-treated patients. CD4^+^ T cells were activated by PHA and IL-2 for 24h while NK cells were pre-incubated with the NaMiX molecules. CD4^+^ T cells were superinfected by spinoculation with HIV III-B and incubated with stimulated NK cells for two or five days at an E:T of 1:1. **A)** P24 intracellular staining of CD4^+^ T cells of one representative donor (left panel) and HIV-1 RNA in the supernatant measured by ddPCR of one representative donor (right panel). The experiment has been performed with three different donors showing the same trend but with high baseline variability. **B)** Human PBMCs were pre-incubated for 48h with the different molecules and stimulated with ACH2 cells during 5 hours. Cells were further stained for extracellular markers to identify CD25 and CD69 expression on CD3^-^CD56^+^CD16^+^ NK cells and CD3^+^CD4^+^ T cells using anti-CD3, CD4, CD14, CD16, CD19 and CD56 antibodies. The figure represents three independent experiments with three different donors. Data were expressed as the mean value ± SD. Statistical analysis was performed using a one-way ANOVA and post-hoc Tukey test (* p < 0.005, **p < 0.005, ***p < 0.0005, ****p < 0.00005) **C)** representative expression of CD25 and CD69 on CD4^+^ T and NK cells pre-activated by NaMiX during 48 hours for one donor included in panel B. **D)** HIV-1 mRNA release in the supernatant of ACH2 cells stimulated during 24 hours by PBMCs prestimulated by NaMiX during 48 hours. Three independent experiments has been performed with three different donors. Data were expressed as the mean value ± SD. Statistical analysis was performed using a one-way ANOVA and post-hoc Tukey test.

### NSG tg-hu-IL-15 humanized mice develop more potent NK cells than NSG mice

We next wanted to evaluate NaMiXs as a potential functional cure in a humanized mouse model of HIV-1 latency. We have already shown that humanized NOD.*Cg-Prkdc^scid^Il2rg^tm1Wjl^*/SzJ (NSG) mice infected with HIV-1 and treated with cART exhibit virological and immunological characteristics similar to HIV infection and HIV latency in humans [36]. However, due to a full IL-2Rγ knock out, human NK cell engraftment and maturation is very low in this mouse strain. Humanized NOD/Shi-*scid*-IL-2Rγ^null^ (NOG) or SIRPA Rag-/-IL2Rγ-/- (SRG) mice transgenic for human IL-15 (tg huIL-15) were developed to have more functional and more mature NK cells [37]. In order to characterize human NK cells, NSG and NSG tg-huIL-15 were humanized with CD34^+^ cells from human umbilical cord blood and sacrificed after four months to evaluate NK cell maturation and cytotoxicity. NSG tg-huIL-15 mice develop ten-times more human NK cells in blood (NSG mean: 2.3% ± 1.2%; NSG tg huIL-15: mean 22.03% ± 13.79%) and lung (NSG mean: 3.8% ± 1.4%; NSG tg huIL-15: mean 36.9% ± 9.6%) and four-times more in spleen (NSG mean: 2.2% ± 0.63%; NSG tg huIL-15: mean 8.9% ± 4.45%) compared to normal NSG mice (Figure 10A-B). Furthermore, NK cells from NSG tg-huIL-15 differentiate into CD56^dim^ and CD56^bright^ subpopulations in blood, lung and spleen (Figure 10C). However, there was only a small increase in overall KIR receptor expression on NK cells from blood (KIR2DL1/DS1 and KIR2DL2/DL3/DS2) and spleen (KIR2DL1/DS1) while NKG2A expression significantly increased in all organs of NSG tg-huIL-15 as compared to NSG mice (Figure 10D), reaching at least 60% of cells expressing NKG2A in the spleen and 80% in the blood. To determine the cytotoxic activity of NK cells in both mouse strains, we incubated splenocytes with K562 cells and stained them for CD107a, intracellular IFN-γ and perforin expression. NK cells from humanized NSG tg-huIL-15 had a greater CD107a-reflected degranulation capacity against K562 cells, accompanied by a significantly higher perforin expression compared to NSG mice (p=0.0186 and p=0.0005 respectively) (Figure 10E). More importantly, splenocytes from NSG tg-huIL-15 were able to kill nearly 70% of target K562 cells (mean 68.75% ± 12.8%) while NSG splenocytes only killed 30% of target cells (mean 29.16% ± 23.61%) (Figure 11C). However, we observed no difference in IFN-γ expression in both mouse strains (Figure 10E). Overall, NSG tg-huIL-15 show a better NK cells engraftment after humanization, and a proper differentiation into fully functional killer cells.

**Figure 10:**
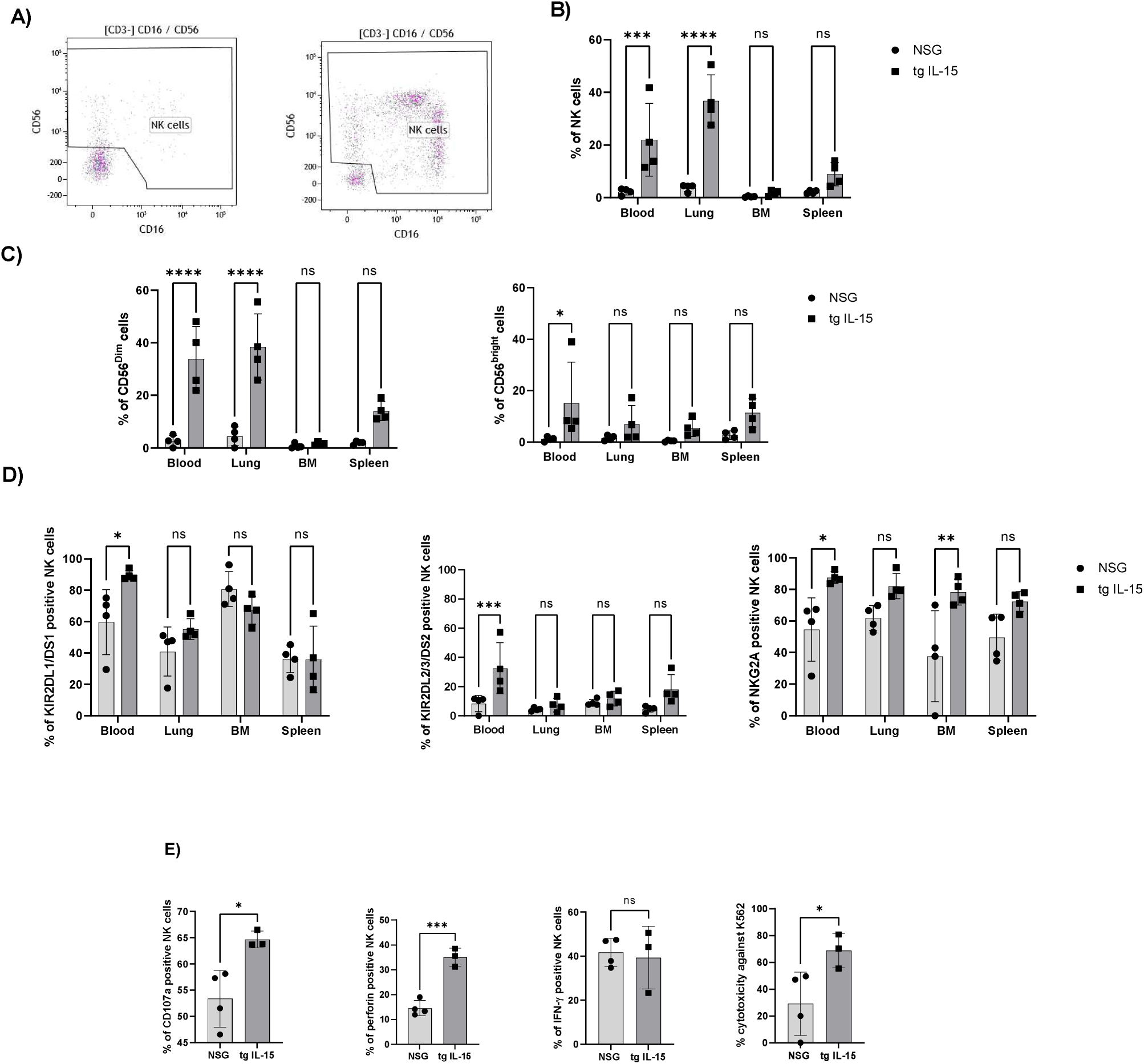
Humanized NSG mice transgenic for human IL-15 develop functional NK cells as compared to NSG mice. NSG mice (n =4) or NSG tg huIL-15 mice (n=4) were humanized with human CD34^+^ cells from umbilical cord blood. After four months, mice were sacrificed and organs harvested. Cells were stained to gate on CD3^-^CD56^+^CD16^+^ NK cells with anti-CD3, CD8, CD14, CD16, CD19 and CD56 antibodies. **A)** Representation of the dot plot of NK cells from the spleen of NSG (left) or NSG tghuIL-15 (right). **B)** Comparison of NK cells in blood, lung, and bone marrow (BM) and spleen in NSG mice and NSG tg huIL-15 mice. **C)** Comparison of NK subpopulations CD56^dim^ (left) and CD56^bright^ (right) in blood, lung, BM and spleen. **D)** Cells from blood, lung, BM and spleen were stained for NK cell markers such as KIR2DL1/DS1 (left) KIR2DL2/3/DS2 (middle) and NKG2A (right). Data were expressed as the mean value ± SD. Statistical analysis was performed using unpaired Student t test. (**p < 0.005, ***p < 0.0005, ****p < 0.00005). **E)** NK cells from Humanized NSG mice transgenic for human IL-15 are cytotoxic as compared to NSG mice. Splenocytes were incubated with K562 cells for 5h together with anti-CD107a (left graph), and were further permeabilized and stained with anti-perforin and anti-IFN-γ and anti-CD3, CD8, CD14, CD16, CD19 and CD56 antibodies to gate on CD3^-^CD56^+^CD16^+^ NK cells. K562 cells were pre-stained with CellTrace Violet and incubated with splenocytes for 5h and all cells were stained for Live/Dead to determine cytotoxicity-induced by NK cells. Data were expressed as the mean value ± SD. Statistical analysis was done using unpaired Student t test. (*p < 0.05, ***p < 0.0005).

### The α.anti-NKG2A.IL-15 NaMiX increased NK cell cytotoxicity *in vivo* but did not stop viral load rebound after treatment interruption

Since α.anti-NKG2A.IL-15 was one of the lead NaMiX molecules to enhance NK cell degranulation and cytotoxicity against target cells *in vitro*, and given the high NKG2A expression of NK cells in humanized NSG tg-huIL-15 mice, we finally evaluated its efficacy *in vivo* in an HIV-1 latency model. 16 humanized mice were infected with HIV-1 for four weeks, treated with cART for six weeks and injected with the molecule or PBS (8 mice per group) ten and three days before treatment interruption. Four mice per group were sacrificed three days after treatment interruption to harvest cells from blood and spleen for phenotyping, cytotoxic activity and degranulation analysis against activated ACH-2 cells. NK cells from the spleen of mice treated with the lead NaMiX had a stronger CD107a expression against activated ACH-2 cells than PBS treated mice (64 ± 8.2 vs. 43.37 ± 9.7%) (p=0.0176) (Figure 12A). In addition, splenocytes from mice treated with α.anti-NKG2A.IL-15 killed 42.38 ± 11.76% of ACH-2 cells whereas NK cells from non-treated mice killed only 23.69 ± 5.5% of ACH-2 cells (p=0.0283) (Figure 12B). We also observed a slight but significant increase in the cell activation marker HLA-DR on blood NK cells from mice treated with NaMiX (mean 9.1 ± 1.09%) as compared to the PBS control group (mean 5.76% ± 1.79) (p=0.0465) (Figure 12C). Even if not significant this tendency was also observed in the bone marrow (p=0.0658). We additionally noticed a lower exhausted CD56^neg^CD16^+^ subpopulation in the bone marrow of α.anti-NKG2A.IL-15 treated mice (mean 77% ± 2.93%) (87.9% ± 5.8) (p=0.0332), which was compensated by a higher functional CD56^dim^ subpopulation (10% ± 3.0 vs. 2.9% ± 2.4%, p=0.0173) (Figure 12D). Interestingly, we did not see a difference in IFN-γ expression on NK cells nor a difference in CD8^+^ T cell activity between the two groups (data not shown).

**Figure 12:**
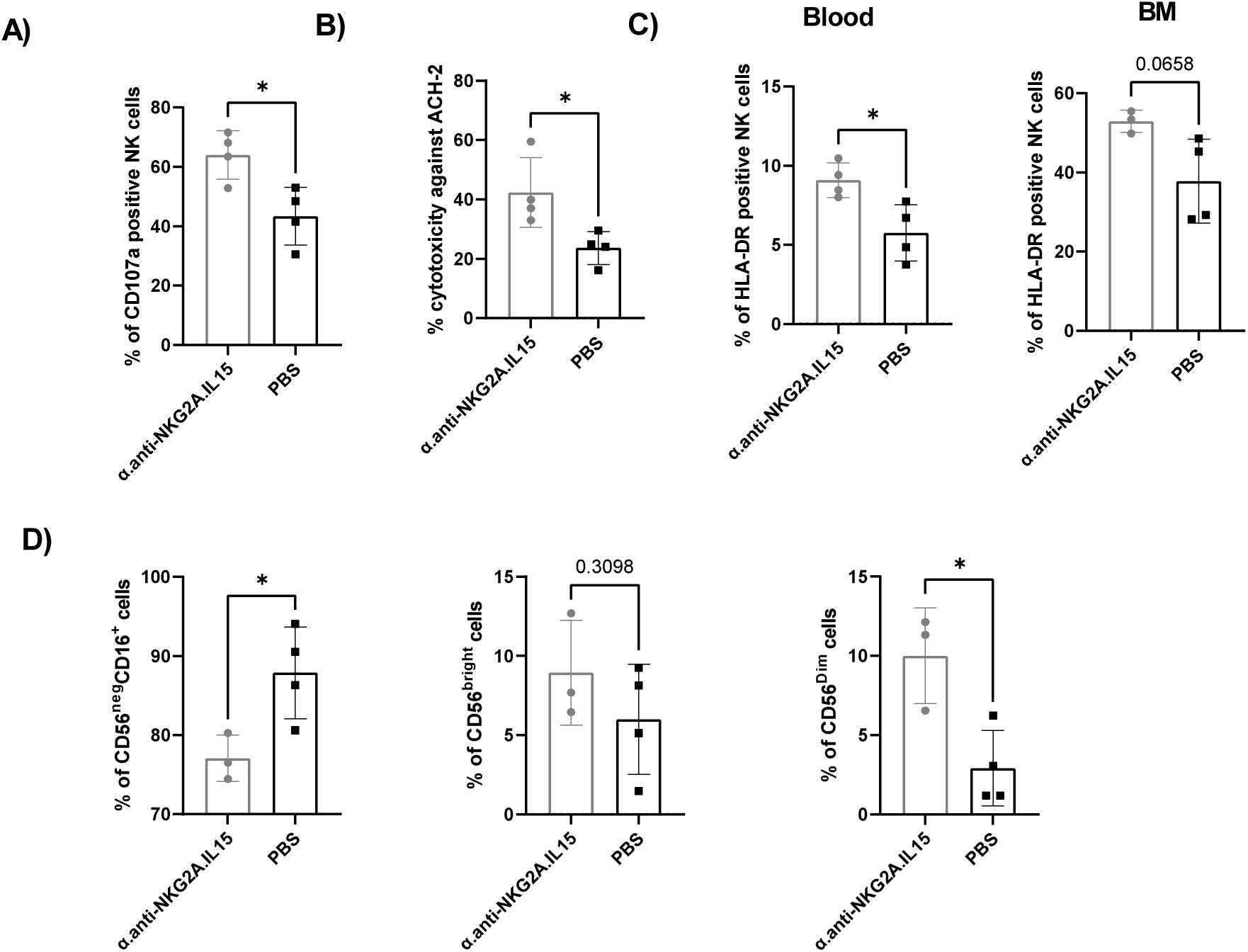
α.anti-NKG2A.IL-15 increased the cytotoxic activity of NK cells in humanized NSG tg huIL-15 mice infected with HIV and treated with ART. Humanized NSG tg huIL-15 mice (n=8) were infected with HIV-1 JRCSF for 4 weeks, treated with cART and injected with PBS (n=4) or α.anti-NKG2A.IL-15 (n=4) ten and three days before treatment interruption. Mice were sacrificed and organs harvested 3 days after treatment interruption. **A)** Splenocytes were incubated with activated ACH-2 cells for 5h together with anti-CD107a and stained with anti-CD3, CD8, CD14, CD16, CD19 and CD56 antibodies to gate on CD3^-^CD56^+^CD16^+^ NK cells **B)** ACH-2 cells were gated and evaluated for Live/Dead staining. **C)** Blood (left) and bone marrow (right) cells were stained to gate CD3^-^CD56^+^CD16^+^ NK cells and the activation marker HLA-DR. **D)** Comparison of NK subpopulation CD56^neg^ (left) and CD56^brigh^ (middle) CD56^dim^ in the bone marrow. Data were expressed as the mean value ± SD. Statistical analysis was performed using unpaired student t test (*p < 0.05).

After cART treatment interruption, we monitored plasma viral load (VL) over several weeks on eight additional mice (four in each group). In this first experiment, cART treatment did not decrease VL to an undetectable level for all mice in both groups (Figure 13A) as it was previously observed in NSG mice [36]. The PBS treated mice showed a heterogeneous VL distribution over time after cART interruption. In contrast, treatment with NaMiX seemed to have homogeneously increased VL in all mice, suggesting a latency reversal effect of the molecule (Figure 13A). In addition, when looking at the overall VL change from the time of cART interruption (week 10) to the end of the experiment (week 14), VL variation was higher in the PBS control group compared to the α.anti-NKG2A.IL-15 treated group (Figure 13B).

**Figure 13:**
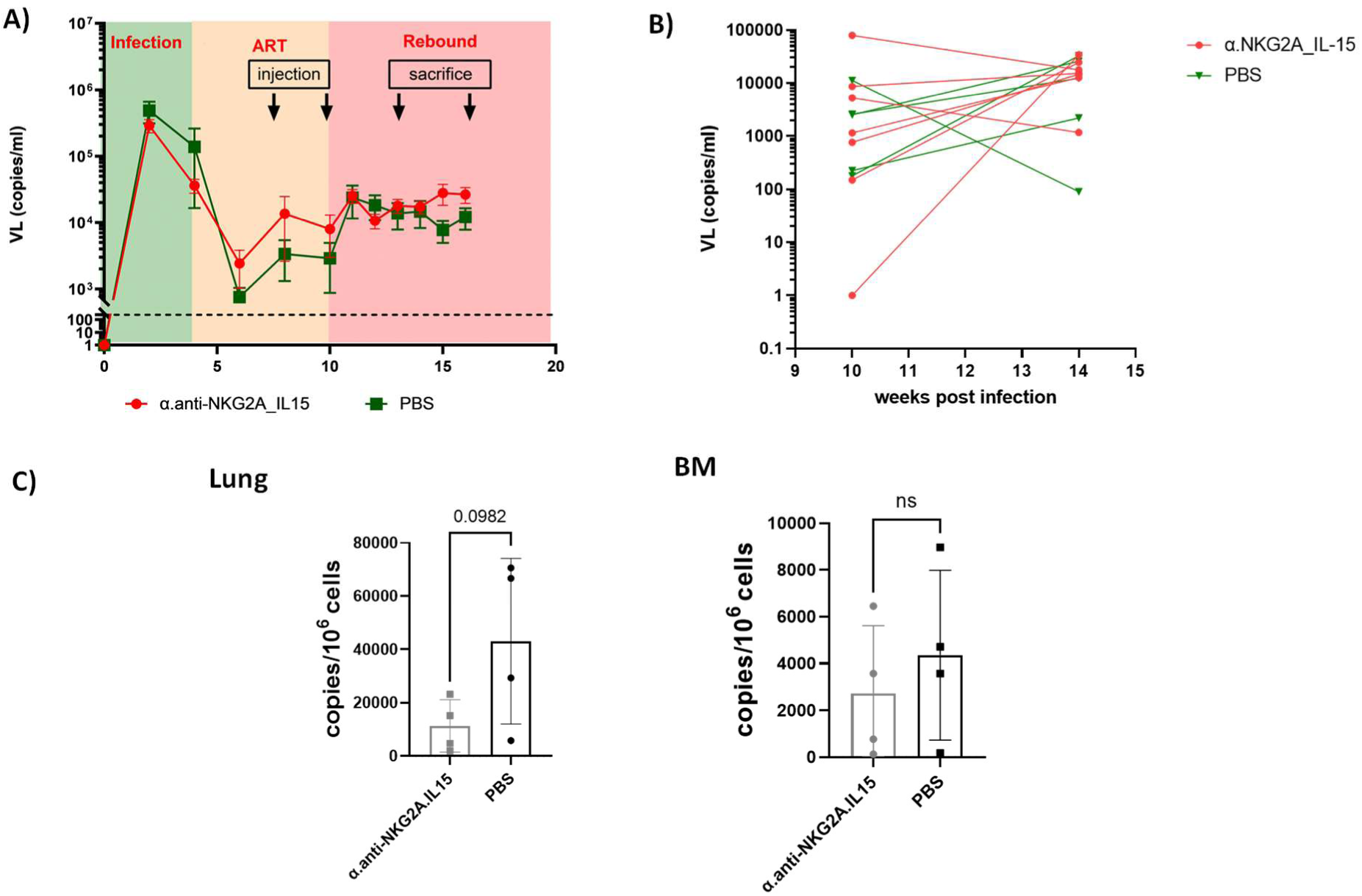
α.anti-NKG2A.IL-15 did not delay viral rebound but decreased HIV-1 reservoir in infected humanized NSG tg huIL-15 mice treated with cART. Humanized NSG tg huIL-15 mice (n = 8) were infected with JRCSF for 4 weeks, treated with cART and injected with PBS (n = 4) or α.anti-NKG2A.IL-15 (n = 4) ten and three days before treatment interruption. Mice were sacrificed and organs harvested 3 days and 7 days after treatment initiation and the last injection of NaMiX (14 weeks after HIV-1 infection). **A)** Viral load (VL) was measured every week or second week. Left panel represents mean VL of the PBS and NaMiX-treated group. **B)** Representation of viral rebound variation between treatment interruption and week 14 for each mice on the right panel. **C)** Total HIV-1 DNA of human CD45^+^ T cells in the lung (left panels) and in the bone marrow (BM, right panel) 3 days after second injection. Data were expressed as the mean value ± SD. Statistical analysis was performed using unpaired student t test (*p < 0.05).

To investigate the role of α.anti-NKG2A.IL-15 as a LRA *in vivo,* we measured total HIV DNA in human CD45^+^ cells in BM and lung three days after the last injection of NaMiX and ART interruption in 4 mice from each group. Even if not significant, we observed a decrease in total HIV-DNA in the lungs of mice treated with the NKG2A NaMiX compared to PBS (p=0.982) (Figure 13C) but not in the bone marrow where immature NK cells reside. These preliminary data indicate that although NK cells were stimulated by NaMiX *in vivo*, there are some evidence that NaMiX could act as a latency reversing agent in these mice. The dose and the time/duration of NaMiX delivery need be further optimized to obtain a significant effect on viral rebound, or to be associated with other therapeutic options to delay viral rebound.

## Discussion

The present work shows that targeted multimerisation of IL-15 on NK cells by the NKG2A or the KIR receptors is a promising approach to enhance its stimulatory functions, and could potentially circumvent the delivery of high and toxic doses of the cytokine. NaMiX increased cytotoxicity and killing activity of NK cells against both cancer *in vitro* and HIV-1 infected cells *in vitro* and *in vivo*. In addition, the molecule can provide further benefits as a latency reversal agent to diminish the size of the viral reservoirs.

Due to the early establishment of the latent reservoirs, researchers have to be very inventive to find a cure for HIV-1 infection. One of the most studied and promising strategies is the “shock and kill” therapy, which relies on the activation of latent reservoir cells by LRAs followed by the recognition and elimination of cells harboring the reactivated virus by NK cells and cytotoxic T lymphocytes (CTL). Purging the entire reservoir without toxicity has proven to be challenging. For example, IL-2 and anti-CD3 antibody stimulation induce a cytokine storm, cytotoxicity and antibody production against anti-CD3 [38], while IL-2 and IFN-γ stimulation has no effect on viral load (VL) rebound [39]. New safer, more specific and potent LRAs have been developed, such as HDACi and HMTi [40–42], NF-кB modulators [43] and TLR agonists [44–46]. Yet, when used alone, none of these LRAs were able to sufficiently decrease the reservoir size nor to delay viral rebound, indicating a deficiency in the “killing” of the reactivated cells. The functions and frequencies of cytotoxic NK and CD8^+^ T cells are highly affected as disease progresses towards a chronic phase, which makes them probably unable to clear the latent reservoir after viral reactivation [10]. Silencing the reservoir, thus avoiding viral rebound, can be achieved by targeted therapeutic vaccines promoting a sustained host immune response against HIV-1. The first vaccine trials aimed to prime CTLs against HIV-1 proteins [47, 48] or to use autologous DCs pulsed with inactive HIV-1 to stimulate CD8^+^ T cells [49, 50]. While these strategies have not demonstrated sustained viral control, cocktails of broadly neutralizing antibodies (bNAbs) are currently the most promising immunotherapeutic approach. Some studies have shown that bNAbs can decrease VL to an undetectable level in infected humanized mice and rhesus monkeys [51], and delay time to viral rebound in human [52]. Furthermore, bNAbs seem to reduce viral reservoirs by stimulating the CTL response [51, 53], and combining latency reversal with bNAb treatment significantly prevented viral rebound after ATI for more than 6 months in half of the infected-treated macaques [54].

NK cells play a key role in the control of viral infection and yet, they were hardly considered as a primary target for any of these cure strategies. Immunotherapies priming NK cells against tumor cells by immune checkpoint inhibition or bispecific and trispecific killer cell engager (BiKEs and TriKEs) just underwent first phase clinical trials [55, 56]. Huot et al. recently described that the non-pathogenic model of African green Monkey, a natural host of SIV, displayed expansion of a terminally differentiated NKG2A^low^CD16^+^ phenotype, while there is accumulation of intermediarily differentiated NKG2A^high^CD16^+^ NK cells in the pathogenic macaque model [57]. We propose here a therapeutic bispecific molecule combining scFvs of the immune checkpoint inhibitors anti-NKG2A or anti-KIR2DL with IL-15, forming dimers or heptamers. At first, we demonstrated that multimerizing IL-15 at the surface of NK cells by targeting NKG2A significantly increased phosphorylation of STAT5 in NK and CD8^+^ T cells compared to IL-15 alone or by targeting KIR2DL. Signal transduction of IL-2Rγ_C_ chain stimulation can either occur through JAK1/3 mediated STAT5 phosphorylation [34] or PI-3-Akt-mTOR mediated phosphorylation of the ribosomal protein S6 [58], inducing survival or proliferation, respectively. It has been proposed that soluble IL-15/IL-15Rα preferentially activates the STAT5 phosphorylation pathway whereas trans-presentation by DCs preferentially activates S6 phosphorylation. We cannot exclude an increased proliferation signal through NaMiXs since high levels of IL-15 also potentially increase S6 phosphorylation [58]. Interestingly, CD8^+^ T cells expressing low levels of NKG2A and KIR2DLs were also stimulated by the STAT5 pathway, suggesting that NaMiX could additionally bind target cells through their IL-15 entity.

We further showed that multimerizing IL-15 combined with NKG2A targeting increased degranulation and IFN-γ expression of NK cells against resistant cancer cells expressing multiple HLAs [59], while KIR2DL targeting did not show a significant effect probably due to reduced receptor expression. The expression of certain NK cell receptors, such as NKG2A and to a lower extent KIR2DL2/3, can be triggered on CD8^+^ T lymphocytes by T cell receptor (TCR) or IL-15 stimulation after a few cell cycles [60]. NKG2A blocking in combination with PD-1 blocking antibodies (monalizumab and durvalumab respectively) was only pertinent when NK and CD8^+^ T cells were pre-stimulated for nine days with IL-15, which increased NKG2A and PD-1 expression [29]. Andre *et al.* further reported that treating NK cells with monalizumab could restore CD107 expression on NKG2A^+^ NK cells against HLA-E positive K562 cells to a similar level seen with wild typel K562 cells. However, the increase in CD107a and IFN-γ was only significant when NK cells were pre-stimulated with IL-2 for 7 days. In contrast, we noticed a significant increase in CD107a and IFN-γ expression on NK cells against HLA-E expressing K562 when incubated with α.anti-NKG2A.IL-15 or β.anti-NKG2A.IL-15, without any pre-stimulation. Andre *et al*. further showed that CD8^+^ T cells primed with flu peptides, stimulated with IL-15 and treated with monalizumab and durvalumab modestly increased CD107a and IFN-γ expression against flu-peptide pulsed K562 [29]. In our study however, CD8^+^ T cells stimulated with NaMiXs did not increase IFN-γ expression when incubated with Raji or HLA-E expressing K562 cells. Similarly, Ohkawa *et al*. showed that cytokine-stimulated CD8^+^ T cells and NK cells, with increased IFN-γ secretion by NK cells, were unable to kill Raji cells [61]. Although blocking NKG2A restored the ability of NK cells to kill Raji cells in their study, we did not observe a significant effect of the β.anti-NKG2A control molecule on the degranulation or cytotoxicity of NK cells and CD8^+^ T cells against Raji targets. We cannot exclude a lower activity of our anti-NKG2A scFv and a potential implication of the immune checkpoint blockade on killing Raji cells when stimulating with the α or anti-KIR2DL forms of NaMiX due to missing controls without IL-15. However, we were able to demonstrate that NaMiXs combining NKG2A blockade with IL-15 stimulation increased cytotoxicity against HLA-E expressing K562 cells suggesting that NaMiXs require both functions to efficiently stimulate NK cells for killing, and that both mechanisms could synergize *in vivo*.

Of note, all molecules were also able to increase degranulation and IFN-γ expression on NK cells and CD8^+^ T cells when exposed to HIV-1 positive ACH-2 cells. In contrast to Raji, HIV-1 positive cells activate the TCR signaling pathway on CD8^+^ T cells, which is the major driver of IFN-γ secretion [62]. Furthermore, β.anti-NKG2A without IL-15 also demonstrated increase in degranulation of NK cells with additional IFN-γ expression on CD8^+^ T cells, indicating that blocking the receptor in the context of viral infection might be more effective than in the context of cancer therapy. We also showed that PBMCs stimulated with all NaMiXs, increase their killing capacity against activated ACH-2 cells to a similar level than seen with 100nM of the IL-15 superagonist [63]. With an estimated molecular weight of 200KDa, this represents 20μg/ml of the superagonist, while we only use 3μg/ml of NaMiXs. The killing seemed to be independent of ADCC, since addition of HIV-1 positive serum did not increase NaMiX cytotoxicity. However, ACH-2 cells do not express CD4 and antibodies from HIV-1 positive individuals required for ADCC, preferentially target the CD4-gp120 bound epitope [64]. Lee *et al.* showed that ADCC was not induced by NK cells against ACH-2 cells, even when incubated with CD4 mimicking compounds [35]. However, when using gp120 coated CEM.NKr-CCR5 target cells or primary T cells from HIV-1 positive individuals, NK cells from HIV-1 negative donors were able to perform ADCC [65]. The effect of the NaMiXs on ADCC in our model must therefore be taken with caution and requires additional studies with other cell lines.

IL-15 has long been known to enhance survival and expansion of exhausted NK and HIV-1 specific CD8^+^ T cells from HIV-positive individuals [66, 67]. Using the viral inhibition assay, we observed an increase in p24 levels and viral RNA after two days incubation with stimulated NK cells. This observation indicates a potential latency reversal effect of the molecules on chronically infected donors. Jones *et al*. demonstrated that IL-15 and its superagonist ALT-803 reverse latency *in vitro* and prime CD8^+^ T cells for activated cell recognition and killing [24]. Similarly, we observed a control of the viral replication when NK cells were pre-treated with the NaMiXs after five days co-incubation with infected CD4^+^ T cells, strongly suggesting that NK cells were primed for killing infected cells. Furthermore, we observed a similar reduction in intracellular p24 levels when NK cells were stimulated with the IL-15 NaMiXs and incubated for five days with infected CD4^+^ T cells than Garrido *et al*. showed with 25ng/ml of rhuIL-15 after 7 days co-incubation [3]. These authors also demonstrated that IL-15 stimulated NK cells were able to detect and clear HIV-1 producing cells after latency reversal, forging our theory that NaMiXs stimulated NK cells clear reactivated cells.

We further evaluated one of the lead molecules, α.anti-NKG2A.IL-15 NaMiX, in a humanized mouse model of HIV-1 latency having functional NK cells. As preliminary data, we found that treating humanized NSG tg huIL-15 mice infected with HIV-1, under cART with α.anti-NKG2A.IL-15, increased the cytotoxic activity of NK cells against HIV positive cells. In a recent study, stimulation with another common γ_C_ chain cytokine, IL-21, in combination with IFN-α delayed viral rebound after analytical treatment interruption (ATI) in rhesus macaques infected with SIV [68]. In contrast, NKG2A NaMiX did not delay VL rebound but seems to affect the size of the viral reservoirs in the lung. A more robust decrease in the size of the reservoir and the stabilization of the viral load could also potentially be achieved by earlier treatment with the molecules, at the start of cART, when NK cells and CD8^+^ T cells are less exhausted and more responsive to cytokine stimulation. In this context, Seay *et al*. demonstrated that the IL-15 superagonist was able to induce viral control up to 21 days in an acute infection model of humanized mice [63], which was mainly driven by NK cell activation.

In the present work, we did not address the implication of CD8^+^ T cells in the maintenance of the reservoir. Cartwright *et al.* demonstrated in an SIV infected rhesus macaque model that CD8^+^ T cell depletion under continuous cART treatment increased the viral load, highlighting the role of CD8^+^ T cells in the control of efficient latency [69]. In this setting, the IL-15 superagonist N-803 has shown to only reverse latency when CD8^+^ T cells were depleted in SIV infected macaques and HIV-1 infected humanized mice under cART [70, 71]. While we did not observe an effect of the lead NaMiX on the VL rebound, we did observe a tendency to decrease the total HIV-1 DNA in the lung but not in the bone marrow containing immature NK cells. This could indicate a latency reversal effect of the molecule *in vivo* and a subsequent eradication of these cells by activated NK cells. This observation is in agreement with the results obtained in the viral inhibition assays with primary HIV-1 infected cells: a first increase of HIV1-mRNA was observed in the supernatant at 3days followed by a decrease at 5 days (Figure 9A). A first activation of CD4^+^ T cells by NaMiX (Figure 9BC) could support an enhanced viral replication followed by their killing by activated NK cells, as observed in ACH2 cells co-cultured with PBMCs stimulated by NaMiX.

To optimize the effect of NaMiXs *in vivo*, further pharmacokinetic and pharmacodynamic studies will be essential, such as half-life, administration route, administration dose and toxicity. We expect an increased half-life of the NaMiXs compared to rhuIL-15 due to the size of the molecule, which should decrease renal elimination. Multiple studies have enlarged the cytokine to increase the t_1/2_ of IL-15 by combining it with its co-receptor IL-15Rα, by attachment to an Ig-Fc domain or by pegylation [72]. However, the only arrangement that showed significant increase in the half-life is ALT-803, which has an Fc domain flanked by IL-15Rα and co-transfected with IL-15 suggesting that NaMiXs have the potential for increasing the longevity of IL-15 *in vivo*. The administration route of IL-15 has also been proven important in clinical trials and might be a limitation factor in our study. The first phase clinical trial with ALT-803 has demonstrated that subcutaneous (SC) injection increased the longevity of the molecule compared to intravenous (IV) injection in human [73]. SC injection also significantly increased NK and CD8^+^ T cell count, maturation and activation in patients with hematologic malignancies that relapsed after allogenic hematopoietic cell transplantation.

In conclusion, we demonstrated in the current work that multimerizing IL-15 and anti-NKG2A at the surface of NK and CD8^+^ T cells increases the function and killing activity of both cell types *in vitro*. We also showed preliminary efficacy of NaMiX on NK cells in an *in vivo* model of HIV-1 infection. Although the *in vivo* dosage of NaMiX has to be optimized or combined with other therapies such as neutralizing antibodies, our results are promising and open doors to new strategies for HIV cure by targeting NK cells.

## Materials and Methods

### Molecular design of NaMiX

The following cDNA constructs were optimized and synthesized (ProteoGenix SAS, Schiltigheim) for expression of the different molecules: human (hu) IL-15Rα-Sushi (UniProt n°Q13261, aa 31-205) - hu C4bp C-terminal β chain (UniProt n°P20851, aa 137-252) - scFv Z199 (humanized anti-NKG2A, Monalizumab, patent n°US20110052606A1) - 5xHis; huIL-15Rα- Sushi-hu C4bp C-terminal α chain (UniProt n°P04003, aa 540-597) - scFv Z199-5xHis; huIL- 15Rα-Sushi-hu C4bp C-terminal β chain - scFv IPH2102 (anti-KIR, Lirilumab, patent n°WO2014/055648Al) - 5xHis; huIL-15Rα-Sushi-hu C4bp C-terminal α chain- scFv IPH2102 (anti-KIR, Lirilumab) - 5xHis. All expression cassettes were cloned between BglII and NotI of the multiple cloning site of a bi-cistronic pEF-IRESpac expression vector. The signal peptide from the tumor necrosis factor receptor superfamily member 16 (UniProt n°P08138) was cloned between EcoRI and BglII. The pcDNA3.1 vector coding for human IL-15 was synthesized by ProteoGenix SAS (Strasbourg, France).

### Cell culture and antibodies

All molecules were generated from stable transfected HEK293F cells (ATCC CRL-1573) cultured with Dulbecco’s modified Eagle’s medium (DMEM) (Gibco, Belgium). PBMCs isolated from healthy donors (Red Cross Luxembourg), Raji (ATCC CCL-86) and ACH-2 (NIH HIV reagent program ARP-349) cells were cultured in Roswell Park Memorial Institute (RPMI) 1640 Medium (Gibco) with 10 mM HEPES (Gibco), 2 mM L-glutamine and 90% non-essential amino acids (Gibco), 10% heat-inactivated Fetal Bovine Serum (FBS, Life Technologies Europe BV, Belgium), 1 U/ml of penicillin, 1 μg/ml of streptomycin (Pen/Strep, Lonza, Belgium). The NK- 92 MI (ATCC CRL-2408) cell line was grown in Alpha-MEM medium (Gibco) with 2mM L-Glutamine, 12.5% FBS and 12.5% horse serum (Gibco). The myeloid leukemia cell line K562 was purchased from the ECACC, the HLA-E expressing K562 cells were a kind gift from Thorbald van Hall (Department of Medical Oncology, Leiden University Medical Center), and were cultured in complete RPMI 1640 supplemented with 10% heat-inactivated FBS (Life Technologies Europe BV), 1 U/ml of penicillin, 1 μg/ml of streptomycin (Pen/Strep, Lonza, Belgium) and 4 mM of L-glutamine (Lonza). HLA-E expressing K562 cells were sorted and maintained in complete RPMI supplemented with 2 µg/mL blasticidin (Sigma-Aldrich, Belgium). All cells were cultured at 37°C with 5% CO_2_. The following antibodies were used for ELISA and flow cytometry: Il-15 Monoclonal antibody (ct2nu): 16.0157.82, PE IL-15 Monoclonal Antibody: MA5-23561, PE-Cy5 CD14 Monoclonal Antibody : 15-0149-42, PE-Cy5 CD19 Monoclonal Antibody: 15-0199-42 from Thermofisher Scientific. Anti-polyHistidine-Peroxidase antibody: A7058, Anti-Mouse IgG-Peroxidase antibody: A9044 from Merck. APC anti-HIS Tag antibody: 362605, PE anti-human CD159a antibody: 375104 from Biolegend. BV711 Mouse Anti-Human CD8: 563677, BUV495 Mouse Anti-Human CD3: 612940, FITC Mouse Anti-Human IFN-y: 552882, BV421 Mouse Anti-Human CD107A: 562623, BUV737 Mouse Anti-Human CD16: 612786, BV786 Mouse Anti-Human CD56: 564058, BV510 Mouse Anti-Human HLA-DR: 563083, PE Mouse Anti-Stat5: 612567 from BD Biosciences. PE CD158a/h (KIR2DL1/DS1) anti-Human antibody: 130-099-209 from Miltenyi Biotec. APC CD158b1/b2j (KIR2DL2/DL3/DS2) Mouse anti-Human antibody: A22333, HIV-1 p24 core antigen KC57: 6604667 from Beckman Coulter and APC HIV-1 (p24) Human Monoclonal Antibody 28B7: MM-0289-APC from Medimabs, 302632 Human Monoclonal Antibody CD25 from Biolegend, 562617 Human Monoclonal antibody CD69 from BD Biosciences, 558616 KI67 Human Monoclonal antibody from BD Biosciences.

### Establishment of stable cell lines for the production of NaMiX

Cells were co-transfected with the bi-cistronic pEF-IRESpac coding for the NaMiX molecules under study and the pcDNA3.1 coding for rhuIL-15. In order to make stable cell lines, HEK293F cells were transfected following the lipofectamine 3000 (ThermoFisher Scientific) manufacturers protocol. 24h prior to transfection, cells were seeded in 2ml FBS-free OptiMEM (Life Technologies Europe BV) in 6 well plates. 4μg of DNA in a 1:1 ratio was transfected with 5µl of lipofectamine and 4μL of reagent. 1ml of complete DMEM medium was added 24h after the transfection. 48h later, cells were transferred in a 10-cm culture dish and cultured in complete DMEM medium supplemented with 5-20 µg/ml of puromycin (InvivoGen) and 100-500 µg/ml geneticine disulfate (G418) (Carl Roth). Clones were expanded in 96 well plates. Supernatant from single-isolated clones were screened using anti-IL-15/anti-His sandwich ELISAs as described below.

### Purification of NaMiX

The clones expressing the highest levels of molecules were slowly expanded from 24 well plates to five chamber Corning® cellSTACKs® (Corning) in DMEM complete medium supplemented with the appropriate selection antibiotics. After 24h the medium was replaced by FBS–free Opti-MEM (Gibco) medium for 48h. Opti-MEM cultured supernatant was collected, cleared by centrifugation, and filtered using 0.22µm PVDF 1L vacuum filter units (GE-Healthcare, VWR). 20mM final imidazole (Sigma-Aldrich) concentration was added to the Opti-MEM supernatant. Molecules were loaded on a Nickel His-Trap™ Excel column (Cytiva) over 48h on a peristaltic pump at a flow rate of 1ml/min (GE-Healthcare, VWR). Using a BioLogic DuoFlow 10 system (Bio-Rad Laboratories NV) the column was washed with 20 mM phosphate buffer with 500 mM sodium chloride (NaCl) at pH7.2. In order to increase detachment of all molecules, the column was left overnight in elution buffer (20 mM phosphate buffer with 500 mM of NaCl and 1M of imidazole at pH7.2). Purified molecules were concentrated on Amicon® Ultra 15, 10KDa MWCO (Millipore-Merck Chemicals NV/SA) and dialyzed against 2×3L of PBS using 10 kDa MWCO Slide-A-lyzer® dialysis cassettes (ThermoFisher Scientific). The final concentration was measured using a NanoDrop™ microvolume spectro-photometer (ThermoFisher Scientific).

### Molecular characterization through ELISA

To select the molecules after purification, different ELISA were performed. Purified molecules or mouse anti-human IL-15 antibodies were coated on a MaxiSorp™ 96-well flat-bottom *ELISA* plate (ThermoFisher Scientific) for 12h and for 72h, respectively (100ng/100uL PBS/well). All incubations with antibodies were done for 1h at 4°C, washed using 1% PBS (Lonza)/BSA (Carl Roth) and blocked with 5% PBS/BSA. Rabbit anti-His-peroxidase was used for detection of the molecules while the IL-15 detection was done in two steps: first with 100ng per well of mouse anti-IL-15 and then with 100ng per well of rabbit anti-mouse IgG conjugated to HRP. Revelation was done with 1x phosphate citrate buffer (Sigma Aldrich) supplemented with chromogen substrate OPD (ThermoFisher Scientific) and H_2_O_2_ (Sigma Aldrich). The reaction was stopped with H_2_SO_4_. Absorbance was read on the POLARstar Omega (BMG Labtech, Belgium) plate reader at 492nm and 630nm.

### Flow cytometry analysis

To determine the binding of the molecules on their respective receptors, the purified molecules were incubated 30 minutes with either PBMCs from Healthy donors (Red Cross Luxembourg) or cell lines expressing the different receptors. NK-92 MI (ATCC CRL-2408) cells were used for molecules recognizing NKG2A while KIR2DL1, KIR2DL2, KIR2DL3 stable HEK293F cell lines were established to evaluate the molecules recognizing the KIR receptors using pcDNA3.1 vectors (OHu24667C, OHu17046C, OHu55562C) (GenScript). The live cells were stained for NK and CD8^+^ T cell surface markers (anti-CD3, anti-CD8, anti-CD56, anti-CD16, anti-CD14 to gate out monocytes, anti-CD19 to gate out B lymphocytes), with anti-His and anti-IL-15 antibodies and live/dead staining) using the LIVE/DEAD Fixable NearIR Dead cell stain kit (Thermo Fisher Scientific) for all flow cytometry analysis. Acquisition was performed on the Fortessa LSR flow cytometer (BD Bioscience, USA) and analyzed with Kaluza (Beckman Coulter, Brea, California, USA). To evaluate intracellular phosphorylation of STAT5, cells were incubated with the molecules for 1, 10, 20 and 40 minutes and stained for extracellular markers on ice. Intracellular staining was performed after permeabilisation with Perm buffer III (BD Phosflow™) following manufacturers protocol. Cells were acquired by the Fortessa LSR flow cytometer or the ImagestreamX (Amnis Corporation, Seattle, WA, USA).

### Measurement of cytotoxicity and killing against cancer or HIV-1 positive target cells

PBMCs from Healthy donors (Red Cross Luxembourg) were incubated for 24 or 48 hours with 3 µg of the molecules or 10ng rhuIL-15 (StemCell, Belgium) in 1ml RPMI complete medium. To measure degranulation and production of cytokines, PBMCs were collected and further incubated with anti-CD107a and Raji cells, or HLA-E expressing K562 or ACH-2 cells at an Effector:Target (E.T) ratio of 10:1 in a 96 V bottom shaped well plate. After 1h incubation, GolgiStop™ and GolgiPlug™ (BD Biosciences) were added for another 4h. Finally, cells were washed and stained with L/D and NK cells and CD8^+^ T cell surface markers (anti-CD3, anti-CD8 to gate in CD8^+^ T lymphocytes, anti-CD14 to gate out monocytes, anti-CD19 to gate out B lymphocytes, and anti-CD56, anti-CD16, to gate in CD3-CD56+CD16+ NK cells, permeabilized following the Cytoperm/Cytofix protocol (BD, Biosciences) and stained with anti-IFN-γ. For measurement of killing, ACH-2 cells, Raji cells and HLA-E expressing K562 cells were pre-stained with CellTrace™ Violet (Invitrogen) following the manufacturer’s protocol. Briefly, target cells were incubated with CellTrace™ Violet for 10 minutes at 37°C. Staining was stopped by adding 10ml complete RPMI and incubating at 37°C for 10 minutes. PBMCs were collected after 48h pre-stimulation with the molecules and further incubated with the pre-stained target cells at an E.T ratio of 10:1 in a 96 V bottom shaped well plate for 5h. In the case of ACH-2 cells, HIV-1 positive or negative sera (diluted 1:1000) was added to evaluate ADCC. After incubation, all cells were stained for L/D staining and acquisition was performed on the Fortessa LSR flow cytometer (BD B ioscience) and analyzed with Kaluza (Beckman Coulter). Cytokine secretion in the supernatant was evaluated by ELISA MAX Deluxe Set Human TNF-α (BioLegend), ELISA MAX Deluxe Set Human IFN-γ (BioLegend), Human Perforin ELISA^BASIC^ kit (HRP) (MabTech), Human Granzyme B ELISA^BASIC^ kit (HRP) (MabTech) following manufacturers instruction. To investigate NaMiX-mediated activation of PBMCS and further killing of ACH2 cells, PBMCs of healthy donors were prestimulated by NaMiX or controls during 48 hours. CD25 and CD69 expression of NK and CD4^+^ T cells were measured by flow cytometry after being incubated with ACH2 cells for 6 hours. HIV-1 replication via the release of HIV-1 mRNA in the supernatant was quantified by ddPCR as previously described [36] after 24 hours of co-culture.

### Viral Inhibition Assay

PBMCs from HIV-1 infected donors on cART (Ethical approval n° 201105/07 from the National Ethics Committee of Luxembourg, each participant signed an informed consent) were thawed and cultured for 24h in complete RPMI medium. CD4^+^ T cells were isolated from the rested PBMC’s using positive selection beads (Miltenyi Biotech) according to the manufacturer’s instructions. After isolation, CD4^+^ T cells were activated for 24 h in RPMI medium with IL-2 (500 IU/ml) and Phytohemagglutinin (PHA) at 5µg/ml. Natural killer cells were isolated using negative selection beads (Miltenyi Biotech) from the unlabeled cell fraction of the previous CD4^+^ T cell selection. After cell isolation, purity of CD4^+^ T cells and NK cells were confirmed by flow cytometry. Natural killer cells were cultured for 24h at a density of 2.10^6 cells/ml in RPMI medium with and without molecules. Activated CD4^+^ T cells were washed twice with RPMI medium and infected with 50 ng/ml HIV III-B lab strain (NIH HIV reagent program ARP-2222) using spinoculation at 1200g for 2h at 25°C. Infected CD4^+^ T cells and NK cells were washed twice with PBS and resuspended in RPMI medium with 50 IU/ml of IL-2 at an E:T ratio of 1:1 for two or five days. Levels of p24 antigen in supernatant were quantified by p24 ELISA (PerkinElmer) according to the manufacturer’s instructions. Levels of intracellular p24 were quantified using flow cytometry. Cells were stained with anti-CD3, anti-CD4, anti-CD56 and LIVE/DEAD Fixable NearIR Dead cell stain kit (Thermo Fisher Scientific). Double p24 intracellular staining was performed after permeabilization using the BD Cytofix/Cytoperm kit (BD, Biosciences) following manufacturer’s instructions with the following p24 antibodies: HIV-1 p24 clone 28B7-APC (Medimabs) and HIV-1 p24 clone KC57-PE (Beckman Coulter). Acquisition was performed on the Fortessa LSR flow cytometer (BD Bioscience, USA). Viral mRNA was measured in the supernatant as previously described [36].

#### NSG mice immune reconstitution, HIV-1 infection and treatment with ART

NSG (NOD/LtSz-scid/IL2Rγnull) (Charles River Laboratory, France) and NSG tg-huIL-15 (Jackson Laboratory, France) were maintained and bred in a specific pathogen free animal facility of the Luxembourg Institute of Health. All experiments on animals were performed with the authorizations from the animal welfare committee of the Luxembourg Institute of Health and the Ministry of Veterinary and Agriculture of Luxembourg (protocol number: DII-2018-21 and DII-2019-02), and complied with the national legislation and guidelines for animal experimentation in Luxembourg. Mice were humanized as we described previously [36] with CD34+ hematopoietic stem cells isolated from human cord blood (CB) using a magnetic activated cell sorting CD34+ progenitor cell isolation kit (Stem Cell Technologies, Belgium). CB was provided by the Cord Blood Bank Central Hospital University (Liège, Belgium) and was collected after obtaining written infomed consent. The protocol (reference 1513) was accepted by the Ethics Committee of the University hospital of Liège (reference B70720072580). Animals that had over 20% of circulating human CD45^+^ cells were infected by two intraperitoneal (IP) injections of the HIV-1 laboratory adapted strain JRCSF ((ARP-2708 HIV-1 Strain JR-CSF Infectious Molecular Clone pYK-JRCSF, NIH HIV reagent program, 10.000 TCID50) within 24 hours. Infection was monitored by viral load measurement every one or two weeks on plasma from blood samples collected by submandibular bleeding. ART treatment was initiated four weeks post-infection and continued for a total of six weeks. Single tablets of dolutegravir/abacavir/lamivudine (Triumeq®, ViiV Healthcare) were crushed and dissolved in Sucralose MediDrop® (Clear H2O) at the therapeutic concentration of 3.40 mg/ml. Drinking solution was refreshed twice per week. Therapeutic molecules or PBS were injected 10 or 3 days previous cART interruption by IV injection at a concentration of 0,2mg/kg. Viral load was determined as described previously by digital droplet PCR [36]. Four mice per group were sacrificed at the end of the treatment and four mice per treatment were sacrificed six weeks after treatment interruption. Blood, spleen, lungs, bone marrow were collected and processed immediately for cell staining, phenotyping, degranulation, and cytotoxic activity as described above. Human CD45+ cells were purified using positive selection beads (Miltenyi Biotech, Germany) and total HIV-1 DNA was measured by PCR in the bone marrow and the lung by ddPCR as previously described [36].

#### Statistical analysis

Statistical analysis was performed using GRAPHPAD PRISM software. Data were expressed as the mean value ± SD. For all *in vitro* experiments, multiple groups receiving the different molecules were compared using one-way ANOVA and post-hoc Tukey tests. For studies in mice, an appropriate sample size (n = 5) was calculated during the study design to obtain groups with a difference of humanization of 10% by taking into account a common standard deviation of 5% using a bilateral student T test based on a 95% confidence level and homogeneous viral load [36].The level of humanization and viremia was randomized between the groups. The two groups were compared using unpaired Student t test. A P-value < 0.05 was considered to be significant.

## Acknowledgements

The following reagents were obtained through the NIH HIV Reagent Program, Division of AIDS, NIAID, NIH: ACH-2 Cells, ARP-349, contributed by Dr. Thomas Folks; Human Immunodeficiency Virus Type 1 IIIB Tat Protein, Recombinant from Escherichia coli, ARP-2222, contributed by DAIDS/NIAID; produced by ABL, Inc; Human Immunodeficiency Virus 1 (HIV-1), Strain JR-CSF Infectious Molecular Clone (pYK-JRCSF), ARP-2708, contributed by Dr. Irvin S. Y. Chen and Dr. Yoshio Koyanagi.

## Competiting interests

The authors declare no conflict of interest. RS, BB, JZ, XD and CSD are the inventors of the filed patent PCT/EP2022/069311 for NaMiX.

